# Deep Learning and 3D Imaging Reveal Whole-Body Alterations in Obesity

**DOI:** 10.1101/2024.08.18.608300

**Authors:** Doris Kaltenecker, Izabela Horvath, Rami Al-Maskari, Zeynep Ilgin Kolabas, Ying Chen, Luciano Hoeher, Mihail Todorov, Saketh Kapoor, Mayar Ali, Florian Kofler, Pauline Morigny, Julia Geppert, Denise Jeridi, Carolina Cigankova, Victor Miro Kolenic, Nilsu Gür, Chenchen Pan, Marie Piraud, Daniel Rueckert, Maria Rohm, Farida Hellal, Markus Elsner, Harsharan Singh Bhatia, Bjorn H. Menze, Stephan Herzig, Johannes Christian Paetzold, Mauricio Berriel Diaz, Ali Ertürk

**Author notes:** These authors contributed equally to this work (Doris Kaltenecker, Izabela Horvath, Rami Al-Maskari, Zeynep Ilgin Kolabas, Ying Chen).

## Abstract

Many diseases, such as obesity, have systemic effects that impact multiple organ systems throughout the body. However, tools for comprehensive, high-resolution analysis of disease-associated changes at the whole-body scale have been lacking. Here, we developed a suite of deep learning-based image analysis algorithms (MouseMapper) and integrated it with tissue clearing and light-sheet microscopy to enable a comprehensive analysis of diseases impacting diverse systems across the mouse body. This approach enables the quantitative analysis of cellular and structural changes across the entire mouse body at unprecedented resolution and scale, including tracking nerves over several centimeters in whole animal bodies. To demonstrate its power, we applied MouseMapper to study nervous and immune systems in high-fat diet induced obesity. We uncovered widespread changes in both immune cell distribution and nerve structures, including alterations in the trigeminal nerve characterized by a reduced number of nerve endings in obese mice. These structural abnormalities were associated with functional deficits of whisker sensing and proteomic changes in the trigeminal ganglion, primarily affecting pathways related to axon growth and the complement system. Additionally, we found heterogeneity in obesity-induced whole-body inflammation across different tissues and organs. Our study demonstrates MouseMapper’s capability to discover and quantify pathological alterations at the whole-body level, offering a powerful approach for investigating the systemic impacts of various diseases.

**Graphical Abstract:** 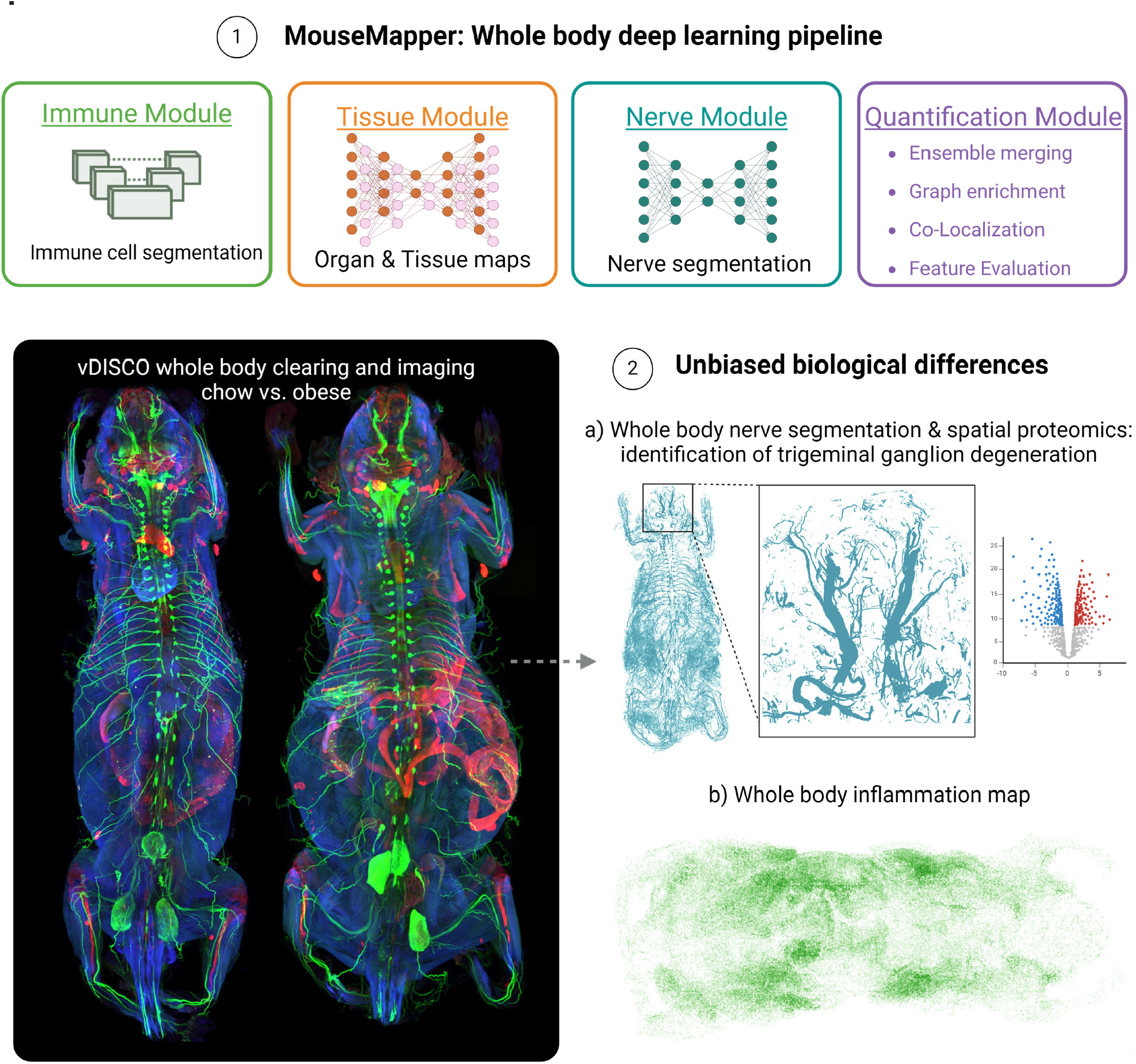

**Highlights:**

1. We developed MouseMapper: AI-driven pipeline for whole-body structural analysis in mice
2. MouseMapper revealed obesity-induced changes in nerves and immune cells across multiple organs
3. MouseMapper identified facial nerve alterations in obesity linked to impaired whisker sensitivity
4. MouseMapper has the potential for holistic 3D analysis of systemic diseases

**Supplementary Videos can be seen at:** http://discotechnologies.org/MouseMapper/

## INTRODUCTION

Many diseases, including lifestyle-induced conditions such as obesity, have far-reaching effects that affect multiple organ systems throughout the body. These systemic effects underscore the interconnected nature of body physiology and the need for holistic approaches to understanding pathological changes. However, tools to study cellular and molecular alterations at the whole-body scale in interconnected systems have been lacking, limiting our ability to understand their broad impacts.

Advanced tissue clearing methods combined with light-sheet fluorescence microscopy (**LSFM**) have enabled the visualization of entire mouse bodies and large human samples at single-cell resolution^1-4^. While these methods permit imaging entire samples such as whole mouse bodies^5,6^, the lack of advanced image analysis tools to quantify cellular and elongated structures such as nerves on a whole-body scale has been a major bottleneck for discovering altered tissue regions in whole bodies. Tools for detecting structural and anatomical changes across scales would also allow selecting regions for further molecular characterization to reveal the mechanisms governing the systemic effects of diseases^7^.

Obesity is associated with chronic low-grade inflammation and a plethora of metabolic dysfunctions such as insulin resistance, impaired glucose tolerance, or hypertension. It increases the risk of developing other comorbidities, including type 2 diabetes, peripheral neuropathies, cardiovascular diseases, stroke and certain types of cancers^8,9^. The connection of obesity to a wide range of diseases underscores the systemic effects that large accumulations of body fat can have on overall health. Recognizing obesity as a systemic challenge emphasizes the need for holistic characterizations of underlying structural and cellular changes that occur with excess body fat.

To this end, here, we developed MouseMapper, an ensemble of deep learning methods to segment and analyze whole body images of the nervous and immune systems and select regions of interest for subsequent molecular analysis (**Fig. 1a**). MouseMapper has three modules for 1) the segmentation of peripheral nerves, 2) the segmentation of CD68+ immune cells, and 3) mapping of segmented structures to organs and tissues across entire mouse bodies. The combination of these models enabled us to study biological structures in their complete spatial and anatomical context. Using MouseMapper, we identified structural alterations in diverse properties of the nerve and immune cell networks with high spatial resolution. Among others, we discovered axonal alterations of trigeminal nerve innervating whiskers in mice, which were associated with behavioral changes and proteomic alterations related to axon degeneration and remodeling. Our whole mouse body scans are available online for the scientific community interested in further exploring HFD-induced changes in the tissues and organs of their interest.

**Figure 1:**
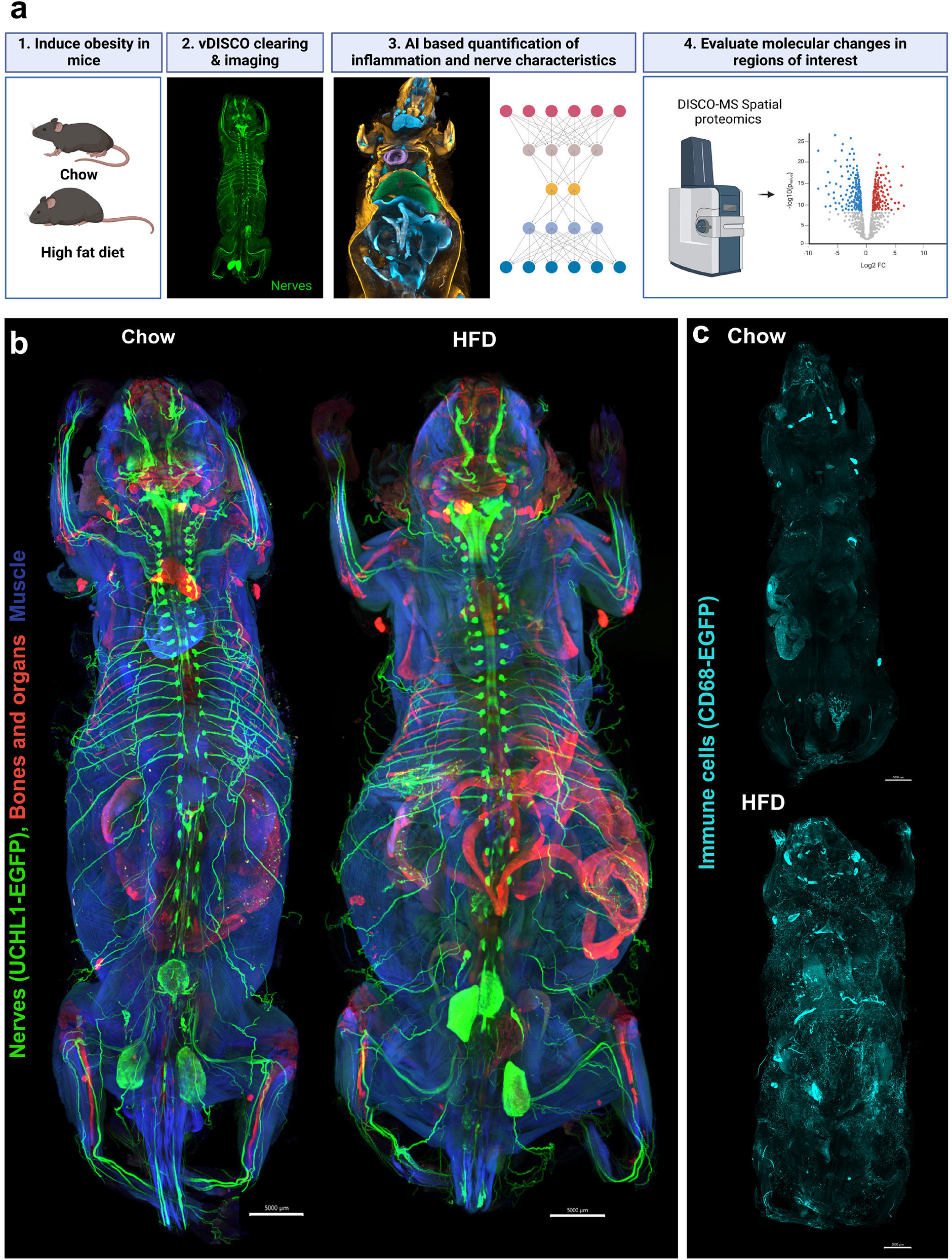
Direct visualization of nerves and macrophages in obesity on the whole-body scale. **a**, Workflow to study obesity-induced changes on the whole-body scale. **b-c**, Representative 3D reconstructions of vDISCO cleared and imaged chow and HFD-fed mice showing b, UCHL1-EGFP+ peripheral nerves and c, CD68-EGFP+ immune cells (n=3/group).

## RESULTS

### Direct visualizations of nerves and immune systems in obesity

In this work we aimed to develop a comprehensive toolset for studying disease-induced whole-body changes, specifically for the study of obesity. To this end, we subjected mice that express EGFP under the peripheral nerve marker UCHL1/PGP9.5 promoter (UCHL1-EGFP mice) or the monocyte/macrophage marker CD68 (CD68-EGFP mice) to HFD-feeding for 16-18 weeks. This led to significantly increased body weights compared to chow-fed controls, mainly due to increased adipose tissue, while lean mass remained similar (**Suppl. Fig. 1a-b**). HFD feeding was associated with impaired insulin response, demonstrating successful induction of metabolic dysfunction in the reporter mice (**Suppl. Fig. 1c-d**). vDISCO clearing and light-sheet imaging of transparent mouse bodies enabled whole-body visualization of the peripheral nervous system (**Fig. 1b, Suppl. Video 1**) and CD68-EGFP+ immune cells (**Fig. 1c, Suppl. Video 2**) in 3D not only in lean mice but also in large obese mouse bodies. For example, in UCHL1-EGFP mice we could clearly trace nerve bundles over lengths of several centimeters, including their paths from the dorsal root ganglia (**DRG**) into the subcutaneous adipose tissue (**ScAT**) depot (**Fig. 1b**). UCHL1-EGFP+ nerves were also visible in internal organs such as heart, liver, spleen, kidneys as well as in brown fat (**Suppl. Fig. 1e-k**). In obese CD68-EGFP mice, an increase in CD68-EGFP+ cell infiltration was apparent throughout the mouse body compared to chow fed controls, with most prominent accumulations in the liver and visceral AT (**ViscAT**) including epididymal, mesenteric, perirenal and cardiac ectopic adipose tissue (**Fig. 1c, Suppl. Fig. 2a, Suppl. Fig. 3, Suppl. Video 3-4**).

### Deep learning ensemble enables whole-body analysis

For an unbiased analysis of obesity induced changes in whole mouse body images, we developed MouseMapper (**Fig. 2a)**, an ensemble of deep learning models that compares animals and conditions. The MouseMapper ensemble consists of three main modules: 1) the Nerve-Module that segments nerves in the entire mouse body and converts them into graphs, facilitating comprehensive mapping and quantitative analysis of nerve characteristics; 2) the Immune-Modul that segments immune cells and quantifies their distributions; and 3) the Tissue-Module that maps organs and tissues to make quantitative data comparable between conditions and animals and to facilitate biological interpretation. The combination of these modules enables a comprehensive description of structural changes in nerves and immune cell distribution across the entire body driven by obesity (**Fig. 2a)**.

**Figure 2:**
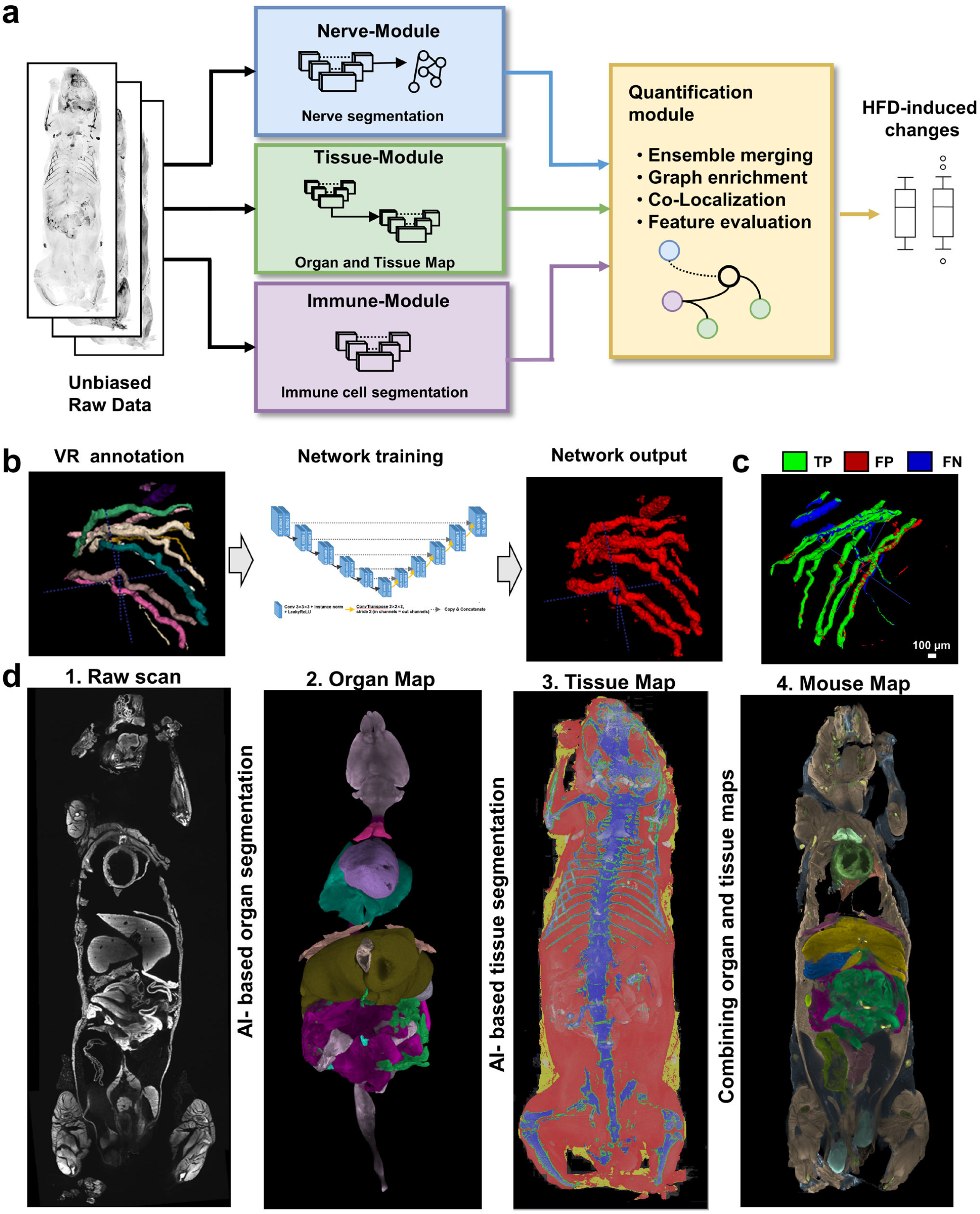
Development of an ensemble of deep learning methods for automated segmentation of nerves, immune cells, organs and tissues. **a**, Workflow: Mice that underwent vDISCO clearing, light-sheet imaging and 3D reconstruction are analyzed using MouseMapper. MouseMapper consists of three modules: the Nerve-Module for deep learning based nerve segmentation, the Immune-Module for deep learning based immune cell detection and the Tissue-Module for automated organ and tissue segmentation. **b**, To train the Nerve-Module, nerves were annotated using virtual reality (VR). **c**, 3D qualitative evaluation of the network performance for the segmentation of nerves based on volumetric dice. Areas that overlap with reference annotations (TP) are masked in green, areas with no overlap in reference annotations (FP) are masked in red. Undetected reference annotation areas (FN) are marked in blue. TP, true positive; FP, false positive; FN, false negative. **d**, The Tissue-Module to segment organs and tissues uses the raw image scan and segments organs (Organ Map) and tissues (Tissue Map), which can be combined to generate a whole mouse map.

Our models were trained in a supervised fashion, with ground truth data being generated using a 3D virtual reality (VR) annotation pipeline^10^. For the Nerve-Module, we annotated nerves in VR (35 sub-volumes with 300×300×300 voxels and 7 sub-volumes with approximately 1000×1000×1000 voxels) from UCHL1-EGFP mouse scans (**Fig. 2b**). We used those patches for training and testing multiple deep leaning models (see Methods). Among the networks tested, the 3D UNet showed the best segmentation performance with a volumetric dice of 0.7913 ± 0.1423 (**Fig. 2c, Suppl. Table 1**) and was used for our nerve segmentation module.

To develop the Immune-Module, we sampled and VR-annotated CD68-EGFP+ cells in five 256×256×256 voxel patches from CD68-EGFP whole mouse scans, representing > 500 contrast-positive cells in adipose tissue and muscle (**Suppl. Fig. 2b**). After training and testing, we found the 3D UNet-based model achieved superior performance compared to other deep learning networks (**Suppl. Table 2, Suppl. Fig. 2c**). In addition, our network was able to segment CD68-EGFP+ cells in other tissues, which were not part of the initial training, including liver and gut (**Suppl. Table 3**) indicating zero-shot inference abilities (transferability to unseen tissue of our model).

Next, we engineered the deep learning-based Tissue-Module (**Fig. 2d, Suppl. Fig. 4a**) that allows mapping segmented structures (e.g., nerves and immune cells here) to organs and tissues, enabling a nuanced interpretation of structural or cellular changes across conditions. To enable efficient organ mapping in the Tissue-Module, given that cell-level accuracy is of lower priority, we down-sampled images of autofluorescence and Propidium iodide (PI) (**Suppl. Fig. 4a**). This enabled the processing of larger volumes by the neural network allowing it to learn shape information while minimizing training time, inference time, and memory requirements. To generate data for training and testing the models, we annotated 20 organs (**Suppl. Table 4)** in each of 10 whole-body mouse scans using VR. We used the annotations from six animals to train multiple neural networks and tested their performance on the other four animals. The 3D UNet architecture implemented in the Tissue-Module again performed best for comprehensive organ segmentation (**Suppl. Table 4**).

While working with down-sampled data provides highly accurate organ segmentation, adipose and muscle tissue segmentation relies on detecting differences in tissue texture, which is not well preserved in the down-sampled images and requires the use of full resolution data. Therefore, we created a separate VR-annotated dataset containing representative patches from muscle, ViscAT, ScAT and brown adipose tissue in full resolution. After completing the training, our network which uses the 3D UNet as the basis showed the best performance compared to other neural networks (**Suppl. Table 5**). By integrating our organ and tissue models, our final Tissue-Module generates a comprehensive anatomical map of the mouse (**Fig. 2d, Suppl. Fig. 4b-f, Suppl. Video 5-6)**. Indeed, the volume extraction of segmented tissue and organs revealed expected increases in adipose tissue (including ViscAT and ScAT) and liver volumes in HFD-fed mice compared to chow-fed mice (**Suppl. Fig. 4g-j, Suppl. Table 6**). In addition, we found that total lymph node mass was increased upon HFD-feeding. Thus, this map serves as a unified reference framework, enabling precise localization of quantitative findings from cellular and anatomical analyses across different body regions.

In summary, MouseMapper represents a robust and automated AI-driven pipeline to detect and quantify system-wide changes in nerve structures or immune cell distributions in any size mouse body.

### AI-based segmentation identifies structural changes associated with behavioral defects in facial nerves in obesity

Obesity is associated with various neuronal malfunctions, including peripheral neuropathies, yet a comprehensive characterization of obesity-induced changes in peripheral nerves on the whole-body scale is lacking. Toward this goal, we applied MouseMapper with the Nerve-Module and Tissue-Module to whole-body scans of normal and obese UCHL1-EGFP mice (**Fig. 1b**). Thereby, we generated whole-body segmentation maps of the peripheral nervous system in both normal and obese mice (**Fig. 3a**). By quantifying the segmented nerve voxels in the whole body and mapping them to specific tissues using the Tissue-Module, we found a significant increase in nerves in adipose tissue, suggesting a concomitant increase of innervation along with increased adipose tissue mass due to HFD-feeding (**Supp. Fig. 5a**). In line, nerve density in adipose tissue was similar between the groups (**Fig. 3b**). Notably, we observed a significant decrease in nerves located in the head (**Fig. 3b**). Most prominently, we observed structural alterations in the infraorbital nerves that innervate the whisker pad upon HFD-induced obesity (**Fig. 3c-d, Suppl. Fig. 5d-f)**. The infraorbital nerve, a key branch of the maxillary nerve and part of the facial trigeminal nerve, is essential for sensory perception, facilitating whisker-mediated tactile exploration and environmental sensing. To quantify the spatial structure of these nerves in more detail, we extracted graphs from the binary nerve segmentation, measuring nerve thickness and the number of nerve endings, but also quantifying the complexity of the nerve network by determining the number and length of edges and the number of nodes/vertices^11^. Here, nodes/vertices represent the points of intersection or branching within the nerve network and edges the connections between these points along the nerve pathways. Quantification of nerve segmentation graphs showed that the number of nerve endings, edges and vertices were reduced by 67.1%, 69.5% and 16.8%, respectively in HFD-fed obese mice (**Fig. 3e-g**). Notably, the thickness of the infraorbital nerve was similar between the chow and HFD-fed mice (**Fig. 3h**), indicating defects in axonal extensions close to the external skin surface rather than a general degeneration of the nerve.

**Figure 3:**
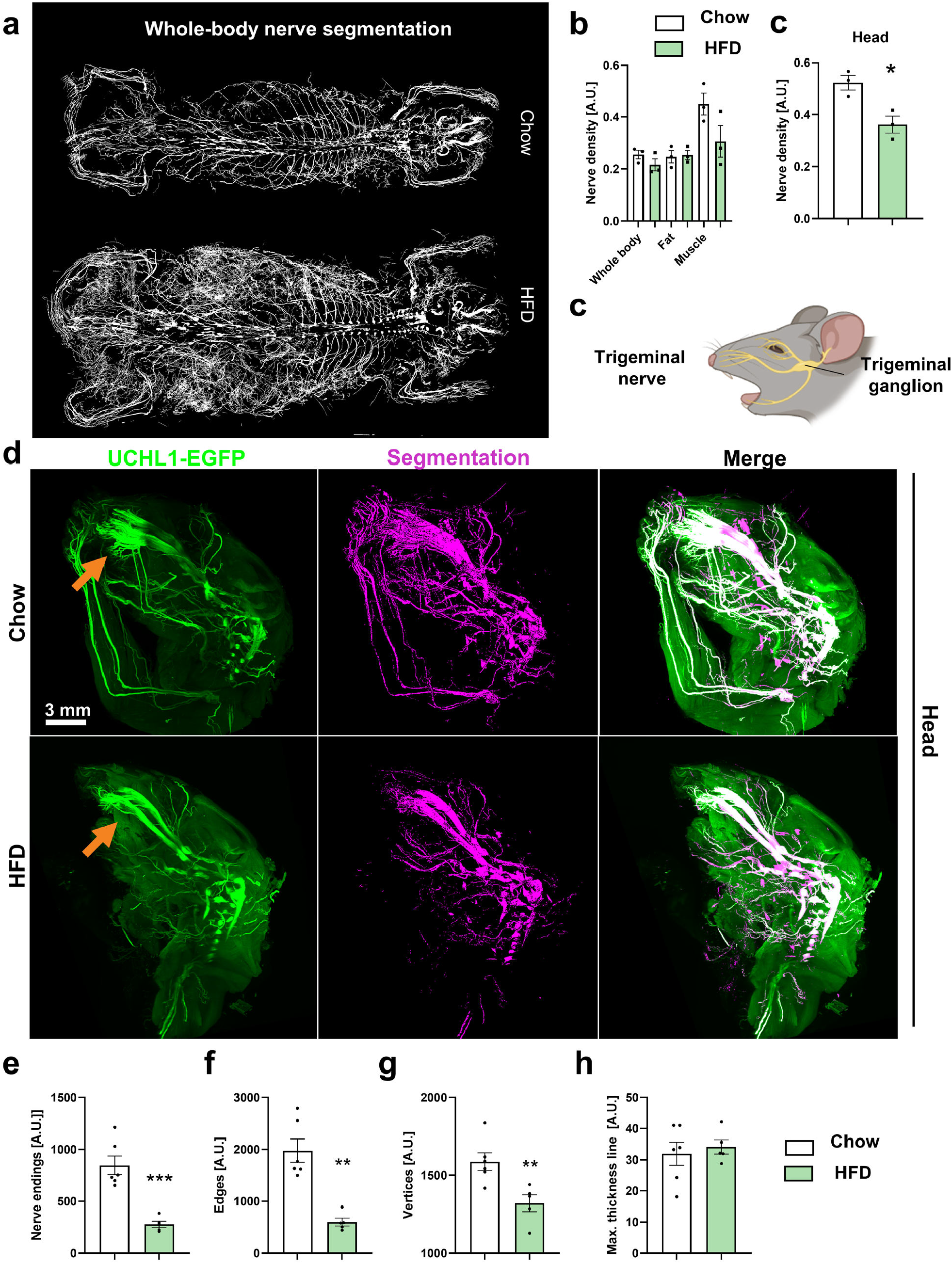
Whole-body nerve segmentation reveal structural changes in the infraorbital nerve in obesity. **a**, Representative whole-body nerve segmentation in chow and HFD-fed mice. **b-c**, Quantification of AI-segmented nerve densities in chow and HFD fed UCHL1-EGFP mice in whole bodies and indicated areas. For the quantification of the nerve density in the head, we masked out the brain for the analysis. n=3/group. **c**, Schematic of a mouse head showing the trigeminal nerve with its three branches arising from the trigeminal ganglion. **d**, Representative heads of chow and HFD-fed mice showing UCHL1-EGFP+ nerves and AI-based segmentation of these nerves. Structural changes in the infraorbital nerve (part of the maxillary branch of the trigeminal nerve) are indicated with the orange arrow. **e-h**, Quantification of characteristics of the infraorbital nerve after graph extractions (n= 6 infraorbital nerves from 3 chow mice, and 5 infraorbital nerves from 3 HFD-fed mice). *p<0.05, **p<0.01.

To assess the functional implications of these structural changes, we performed whisker stimulation tests and found that obese mice exhibited a diminished response to whisker stimulation (**Fig. 4a**). This finding suggests that obesity-induced structural changes in facial nerves may contribute to sensory dysfunction, highlighting the potential importance of our observations.

**Figure 4:**
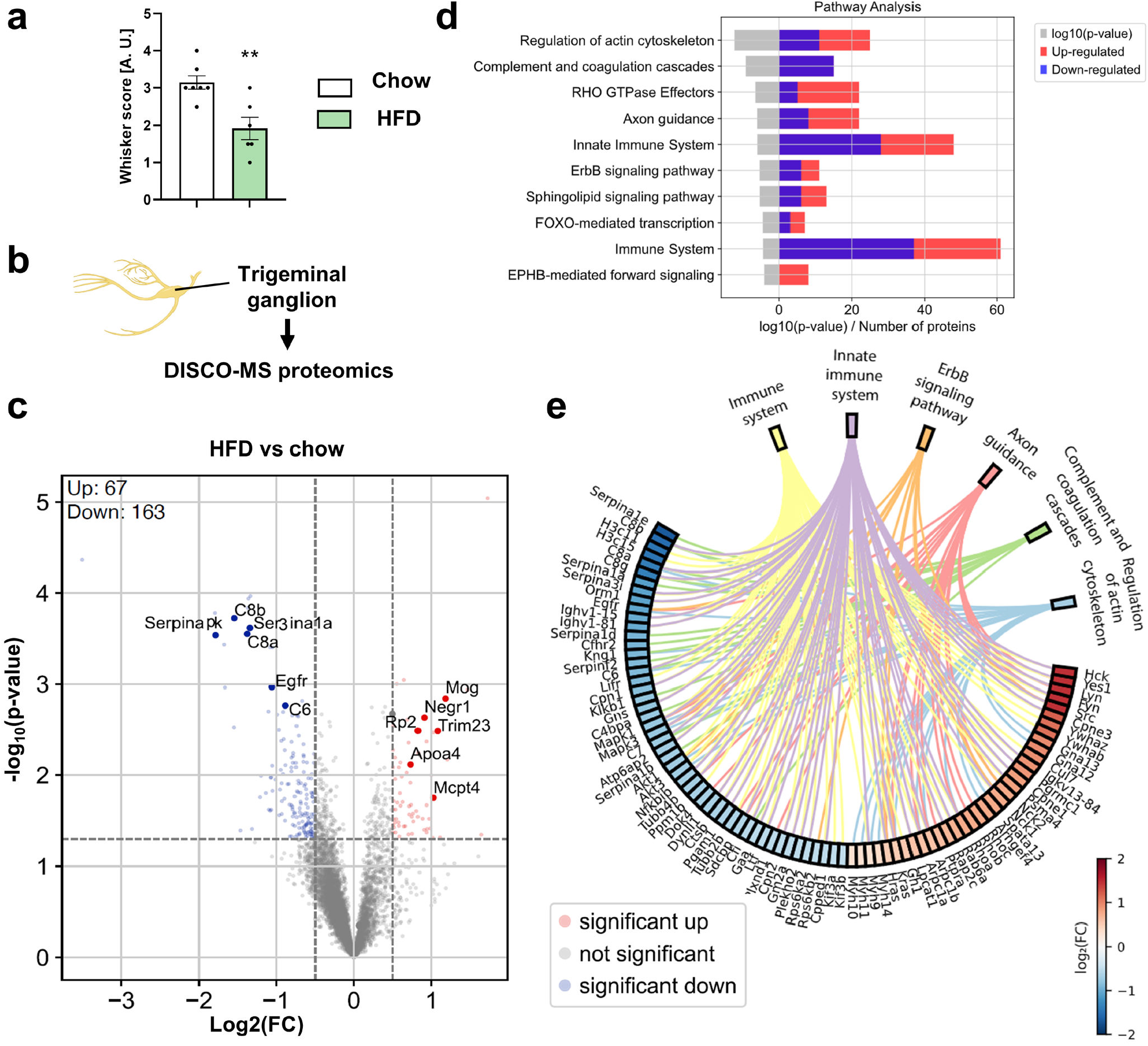
Structural changes in the infraorbital nerve associate with changes in the TG proteome. **a**, Functional assessment of the response after whisker stimulation (n≥6/group, **p< 0.01). **b**, Samples for spatial proteomics profiling were extracted from the trigeminal ganglions of UCHL1-EGFP mice. **c**, Volcano plot showing differentially regulated proteins in chow vs HFD-fed mice. **d**, Pathway analysis showing differentially regulated pathways. Gray bar on the left side of the plot represents the log10 of p-value of each pathway whereas the right side of the plot depicts the number of proteins significantly different in each pathway, red representing the number of up-regulated proteins and blue representing the number of down-regulated proteins. **e**, Chord plot showing a subset of differentially regulated pathways and the corresponding proteins.

### Structural changes in infraorbital nerve coincide with proteomic alterations in the trigeminal ganglion

By developing and applying MouseMapper, we identified and characterized the specification of neuronal abnormalities, particularly in the infraorbital nerve projections of obese mice. Next, to investigate the molecular mechanisms underlying these changes in the infraorbital nerve, we performed spatial proteomics profiling of the trigeminal ganglia, the origin of the infraorbital nerve and the location of their neuronal cell bodies. We dissected the trigeminal ganglia of chow and HFD-fed mice that we imaged above, collected 18G-needle punch-sized samples, and analyzed them using the mass spectrometry-based proteomics (**Fig. 4b, Suppl. Fig. 6a**). We identified more than 6.000 total proteins in each sample **(Suppl. Fig. 6b**). Among them, 230 were differentially regulated (67 up-regulated, 163 down-regulated, **Fig. 4c**) in the trigeminal ganglia between chow and HFD-fed mice.

Pathway analysis revealed multiple differentially regulated pathways in obese mice, including “Regulation of actin cytoskeleton,” “Rho GTPase effectors,” and “Axon guidance” (**Fig. 4d,e, Suppl. Fig. 6c**). This could indicate disruptions in actin dynamics, which is essential for maintaining axonal structure and function. In addition, significantly regulated pathways included “Complement and coagulation cascade”, “Erb-Signalling” and “Sphingolipid signaling” pathways that are involved in inflammation and cellular stress response, among other processes, which also fits to the dysregulation of many general and innate immune pathways (**Fig. 4d,e, Suppl. Fig. 6c)**.

Among down-regulated proteins, we identified multiple members of the serpin serine protease inhibitor (SERPIN) A family (**Fig. 4e**). SERPINA1 has anti-inflammatory properties, especially linked to neutrophils (by inhibiting neutrophil elastase) and is known to protect from tissue damage^12^. SERPINA3 is an inhibitor of cathepsin G, another protease important in neutrophil related immune responses and implicated in inflammation related tissue damage^13^. Downregulation of SERPINA proteins could lead to a reduced ability to control inflammation-induced tissue damage and degradation of structural proteins essential for nerve integrity. We used western blotting for some differentially regulated proteins to validate our proteomic findings in protein lysates of trigeminal ganglion tissue. Thereby, we could confirm the decreased expression of SERPINA1, ERK activation, and increased expression of the GTPase SEPTIN7 (**Suppl. Fig. 6d and e**).

Together, spatial molecular profiling of the trigeminal ganglion in HFD-fed mice, revealed by our MouseMapper deep learning ensemble, showed significant proteomic alterations. Among these, we identified dysregulated pathways related to axon growth and remodeling, and inflammation, which could explain structural changes in the infraorbital nerves.

### Revealing whole-body wide inflammation in obesity

Chronic inflammation is a major hallmark of obesity, intricately linked to the development of various chronic diseases throughout the body. The systemic nature of obesity-induced inflammation underscores the critical importance of understanding which tissues and organs are affected in obese animals and to what degree. To study the spatial context of inflammation in obesity, we applied MouseMapper using the Immune-Module and Tissue-Module to whole-body scans of lean and obese CD68-EGFP mice (**Fig. 5a-b**).

**Figure 5:**
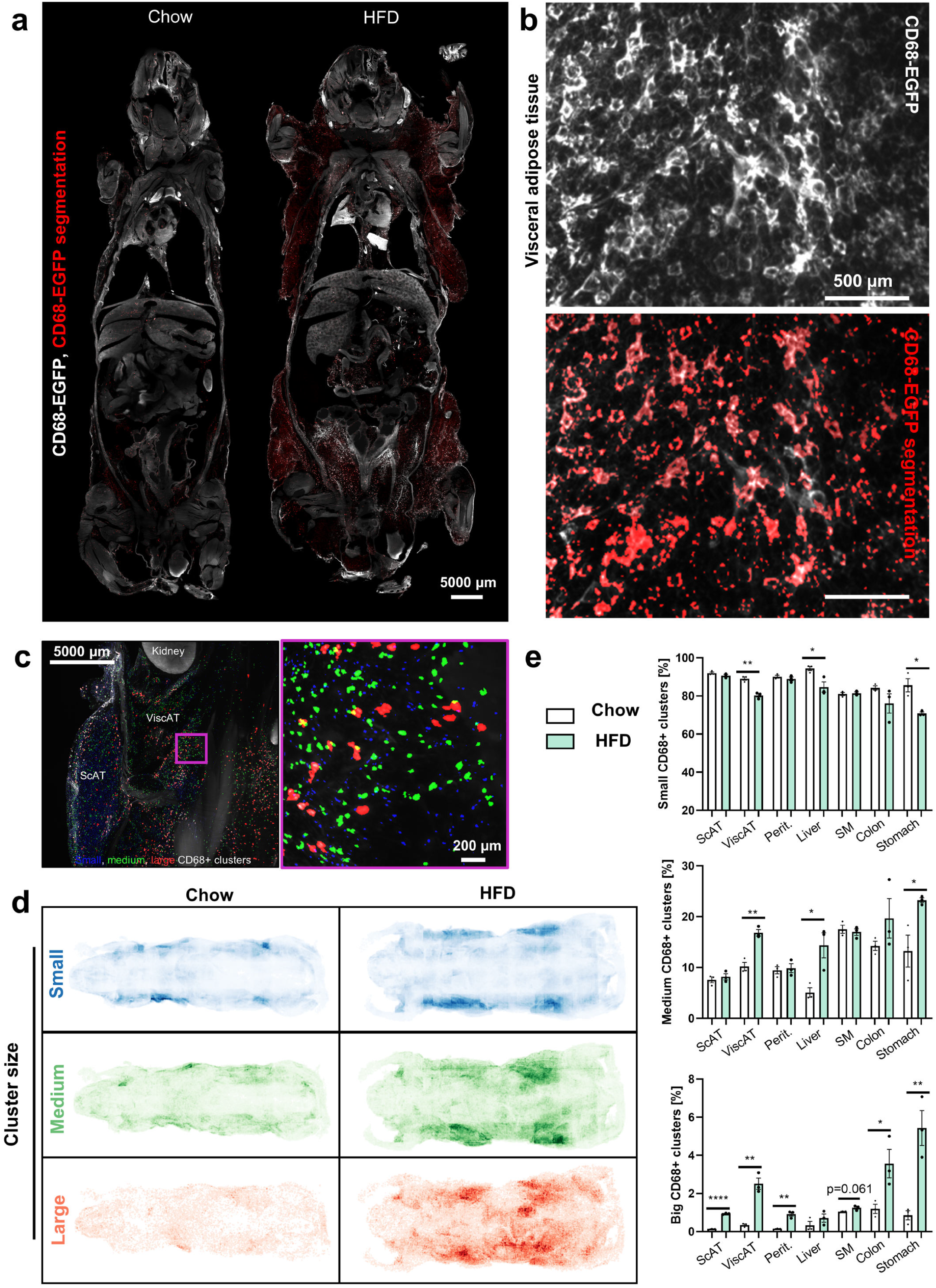
Whole-body wide inflammation in obesity. **a**, Representative images of CD68-EGFP mice (300 μm Z-projection) with AI-based segmentation of CD68-EGFP signal of medium and large sized clusteres overlaid in red (n=3/group). **b**, Z-projection (300 μm) showing ViscAT with CD68-EGFP (upper panel) and AI-based segmentation (lower panel) of an obese mouse. **c**, Segmented cells were grouped into three size clusters: small (blue), medium (green), large (red). **d**, Whole-body CD68-EGFP segmentation results in representative chow and HFD-fed mice showing small, medium and large cluster densities. **e**, Quantification of small, medium and large cluster proportions in indicated organs and tissues. n=3/group, *p<0.05, **p<0.01.

The CD68-EGFP+ immune cells were visible as round, cluster-like structures in tissues such as adipose tissue, liver, skeletal muscle and the peritoneum (**Suppl. Fig. 2a, Suppl. Fig. 3, Suppl. Video 4**). The size of immune cell clusters can indicate the inflammatory state within tissues, with larger clusters correlating with a more activated and pro-inflammatory state^14^. Thus, we generated inflammation maps of CD68-EGFP+ immune cells and grouped them into three different sizes of clusters: small clusters with few cells, medium sized with tens of immune cells and large ones with hundreds of immune cells (**Fig. 5c-d, Suppl. Video 7)**. Using these categories, we analyzed the density of the three different classes of CD68-EGFP+ clusters (**Fig. 5e**) and found notable decreases in small cluster portions within the liver, ViscAT and stomach upon HFD-feeding, whereas this category remained unchanged in ScAT, peritoneum, and muscle **(Fig. 5e**). Conversely, the portion of medium-sized clusters showed an increase specifically in the liver, ViscAT and stomach highlighting a shift from small to medium clusters in these tissues. Additionally, we observed significant increases in large clusters in ScAT, ViscAT, peritoneum, colon and stomach signifying an intensification of inflammatory activity and immune cell involvement in these tissues in obesity (**Fig. 5e**).

Direct visualization of CD68-EGFP+ cells revealed widespread increases throughout the body following HFD-induced obesity. Using the Tissue-Module combined with the AI-based macrophage detection of the Immune-Module, we quantified shifts in CD68+ cluster sizes, confirming elevated inflammatory states across tissues in response to a HFD-induced obesity and providing detailed spatial information.

## DISCUSSION

In In this study, we developed MouseMapper, a deep learning ensemble for comprehensive end-to-end analysis of whole-body systems. Our approach enables 3D organ and tissue mapping of structural changes to study disease-induced changes in biological systems down to cellular resolutions at the whole-body scale without pre-defining specific tissue regions. MouseMapper can faithfully segment elongated nerve structures over centimeters in whole mouse bodies. It can also identify and analyze immune cells from single cells to clusters of hundreds of cells in 3D.

A key strength of MouseMapper lies in its powerful deep learning ensemble, trained on datasets coming from cell-level imaging of whole mouse bodies. This includes nerves traced over long distances using virtual reality in entire mammalian bodies at high resolution - a dataset that presents unique challenges due to the diverse tissue backgrounds encountered across the body, from muscle to bones to various organs. This unique training data enabled us to train 3D UNet models capable of faithfully segmenting complex structures across heterogeneous environments. MouseMapper is built as a versatile, multi-modular system that is easily adaptable to other tissue structures such as blood vessels, lymph vessels, or different types of cellular clusters. Notably, our Tissue-Module provides crucial anatomical context that can localize identified changes within specific organs and tissues and can serve as a common reference framework for other whole-body data. To promote further research and development in this field, we are making our data and algorithms publicly available.

Our key biological findings include structural changes in the infraorbital nerve of obese mice, characterized by reduced axonal extensions and altered network complexity. The infraorbital nerve belongs to the facial trigeminal nerve, which consists of three branches that convey sensory signals from the face through the trigeminal ganglion^15^. Our data reveal a previously unrecognized impact of obesity on facial nerve structure. The reductions in nerve endings and network complexity, suggest a potential mechanism for sensory alterations in obesity, including the reduced sensitivity to whisker stimulation observed by us and the aberrant sensory and pain processing previously observed in obese mice^16-18^. The proteomic changes identified in the trigeminal ganglion offer insights into the molecular underpinnings of these neuronal changes. In this regard, the observed changes in pathways related to cytoskeletal regulation and axon guidance in the trigeminal ganglion could potentially explain the observed changes in infraorbital nerve structure, as both are essential for structural plasticity^19,20^. The proteomic changes related to inflammation underscore the link between obesity and neuroinflammation^21^. These insights could pave the way for novel therapeutic approaches targeting neuroinflammation and cytoskeletal integrity in obesity and related conditions. Notably, these changes likely reflect a combination of neuronal and non-neuronal responses, as neurons comprise only a fraction of cells in the ganglia^22^.

Our data using CD68-EGFP mice support previous findings that obesity is associated with chronic inflammation^23^, as we observed increased expression of CD68-EGFP+ cells throughout the mouse. In line with previous reports, our data confirm a more pronounced accumulation of large CD68-EGFP+ clusters in visceral fat compared to subcutaneous fat^24^. Our whole-body mapping approach adds a comprehensive spatial view of obesity-induced inflammation, revealing tissue-specific patterns of macrophage accumulations.

We made the whole mouse body maps available online, where scientists can easily scroll through large datasets of HFD versus chow-fed mice to investigate neuronal and immune cell alterations (Chow-Nerve, HFD-Nerve, Chow-Inflammation, HFD-Inflammation). Researchers can quickly identify obesity-induced changes in their tissues/organs of interest and explore potential connections with other body systems. These online maps can save time and resources and provide a broader context for understanding localized changes within the global landscape of obesity-induced alterations.

In conclusion, we developed MouseMapper, an ensemble of deep learning tools for characterizing structural changes in whole body systems in response to diseases or other perturbations. We revealed alterations in axons innervating the face in obesity and showed global inflammation associated with regional clustering of immune cells in various areas. While we applied MouseMapper to obesity here, the pipeline can be easily adapted to other complex diseases and other body-wide systems such as the lymphatic and vascular system. In combination with spatial proteomics analysis of hotspots of structural alterations, MouseMapper facilitates the identification of potential therapeutic targets to reverse or prevent pathological changes. MouseMapper thus provides a blueprint for the holistic analysis of complex biological phenomena in 3D.

## MATERIAL AND METHODS

### Animals

8-week-old male UCHL1-EGFP and CD68–EGFP mice on a C57BL/6J background were fed either a chow diet or a high-fat diet (60% fat, #D12492i from Research Diets Inc.) for 16-18 weeks ad libitum. Mice were maintained on a 12-h light–dark cycle. Body composition was determined using an EchoMRI-100H system (EchoMRI, Houston, TX, USA). For insulin tolerance tests (ITTs), mice were fasted for 6h and i.p. injected with 0.75 U/kg insulin. Blood glucose was measured from the tail vein at indicated time points using glucose test stripes. Mice were sacrificed following deep anesthesia with a mix of ketamine/xylazine, followed by intracardiac perfusion with heparinized PBS (10 U/ml heparin) and by a perfusion with 4% paraformaldehyde (PFA). Mice were post-fixed overnight in 4% PFA and subsequently washed five times with PBS shaking (300 rpm) at room temperature for 1h for each wash step. Animal experimentation was performed in accordance with the European Union directives and the German animal welfare act (Tierschutzgesetz). They have been approved by the state ethics committee and the government of Upper Bavaria (ROB-55.2-2532. Vet_02-21-133, ROB-55.2-2532.Vet_02-16-117, ROB-55.2-2532.Vet_02-17-49).

### Whisker stimulation test

The whisker test paradigm was adapted from the methods described previously^25-28^ and the Neuroscore test^29^. To avoid introducing confounding variables, mice were kept in their original cages. A q-tip with a wooden end was used to administer the test. Initially, the q-tip was presented in front of the mouse’s head and allowed to touch it. This was followed by four consecutive strokes, first to the whiskers on the right side and then on the left side of the face. The response to the q-tip stimulation was evaluated using a modified whisker score test. A normal behavioral response to the stimulation, such as turning the head towards or away from the q-tip or initiating grooming, was assigned a score of one. A lack of response to the stimulation was assigned a score of zero. Both sides of the face were stimulated four times, and the scores were recorded by a blinded evaluator. The maximum whisker score was 8, in which mice would have responded to all stimuli. The total score was then averaged for both sides. High scores (3-4) indicated normal responses to the stimulation, while low scores (0-2) suggested a lack of reaction, consistent with sensory deficits.

### vDISCO nanobody labeling and clearing

vDISCO was performed as previously described^2,30^ in combination with GFP-Nanobooster labeling (Atto647N-conjugated anti-GFP nanobooster Chromotek Cat.# gba647n-100;RRID:AB_2629215) for 6 days and passive labeling for 3 days. Mice underwent DISCO clearing^31^ using a Tetrahydrofuran (THF) /H20 series (50% THF, 70% THF x2, 90%THF, 100%THF) for 24h per step followed by an incubation in dichloromethane (DCM) for 6h. Tissues were incubated in benzyl alcohol/benzyl benzoate (BABB, 1:2 v/v) until tissue transparency was reached (>48 h).

### Light sheet fluorescence microscopy

Light-sheet imaging for whole mouse bodies was conducted using a dipping 1.1× objective lens (LaVision BioTec) on an Ultramicroscope Blaze (LaVision BioTec). Tiling scans were acquired with 35% overlap, 100% sheet-width 0.035 NA, 100ms exposure and a 6 μm Z-step size. The images were taken in 16bit depth and at a nominal resolution of 5.9 μm/voxel on the XY axes. In z-dimension we took images in 6μm steps using two-sided illumination. Stitching of tile scans was carried out using Fiji’s stitching plugin with the “Stitch Sequence of Grids of Images” feature^32^ and custom Python scripts.

### 3D reconstruction

Dorsal and ventral scans were fused as previously described^2^ using Arivis and exported whole-body TIFF stacks were used for image analysis.

### Annotation of data in virtual reality for ground truth data generation

Annotations of ground-truth data was performed in virtual reality^10^ using the syGlass software as previously described.

VR-annotation of peripheral nerves was conducted on 35 300×300×300 voxel sub volumes from UCHL1-EGFP mouse scans. Additionally, seven complete trigeminal nerve images of varying sizes were cropped from the UCHL1 channel and included in the annotation. VR-annotation for CD68-EGFP+ cells was performed in five 256×256×256 voxel patches from CD68-EGFP whole mouse scans, selected from representative regions of interest. Annotations were based on both the autofluorescence and CD68-EGFP signal channels. These patches were further cropped down into 40 128×128×128 voxels patches, that were used to train 3D networks for the segmentation of the markers of interest. For the development of the Tissue-Module, we annotated 20 organs of interest in ten downsampled (10-fold) mouse scans (six from CD68-EGFP and four from UCHL1 mice, with five chow-fed and five HFD-fed mice from each line) using the autofluorescence and propidium iodide (PI) channels with the syGlass software. This approach was sufficient to distinguish all organs of interest. To generate reference annotations for the tissue segmentation, we annotated an initial dataset of three 1024×1024×1024 voxels sized patches in full resolution, containing 500 million voxels of fat (visceral, subcutaneous and brown), 145 million voxels of muscle, 16 million voxels of bone tissue, and 8 million voxels of bone marrow. We iteratively increase the size of our annotated dataset through inference on un-annotated patches, and manual correction of the wrongly segmented areas.

### Deep learning-based segmentation for peripheral nerves (Nerve-Module)

To train the peripheral nerve segmentation network, we divided the 35 annotated patches from various parts throughout the mouse body into 28 for training and seven for testing. Similarly, the seven trigeminal nerve images were split into five for training and two for testing. To fit the gpu memory constraints, the five training samples of the trigeminal nerve were further cropped into patches matching the size of the patches from other parts of the mouse body. Consequently, a total of 565 300×300×300 voxel patches were obtained for network training. For model evaluation, we used seven testing patches from various parts throughout the mouse body along with the remaining two trigeminal nerve samples.

Initially, we implemented and trained the following baseline architectures: Attention UNET, NNFormer, SwinUNETR, UNETR, VNet, and 3D UNet. We trained these with a patch size of 128×128×128 voxels, initial learning rate of 1e-3, SGD Optimizer, learning rate decay and Binary Cross Entropy + DICE loss for 1000 epochs.

Upon inspecting the training curves, we observed that the models were almost but not fully converged, and, due to its superior performance, we chose to further train the 3D UNet Architecture.

The final model for peripheral nerve segmentation was trained using the nnUNet pipeline, with a patch size of 128×128×128 voxels, an initial learning rate of 1e-3 with learning rate decay, and the SDG optimizer for 2000 epochs. Additionally, we incorporated the clDICE loss function^33^ into the baseline by adding it with a weight of 0.5 to the original loss. The clDICE loss function aids the model in capturing the topology and connectivity of nerves, leading to more complete nerve segmentation performance.

Before inputting the patches into the network for training or testing, we performed sample-wise percentile normalization. Specifically, for patches from the whole body and every trigeminal sample, we computed the 0.5th percentile and 99.5th percentile of all voxel intensity values to set the minimum and maximum thresholds. Intensity values below the lower percentile or above the upper percentile were clipped to the minimum or maximum thresholds. Finally, we applied min-max normalization. This normalization step enhanced patch contrast by stretching the intensity range between the chosen percentiles and removing outliers, thereby emphasizing nerve regions to improve model performance.

### Deep learning-based segmentation of CD68-EGFP+ cells (Immune-Module)

For training the CD68 segmentation network, we implemented the following architectures: 3D UNet^34^, V-NET^35^, Attention UNET^36^, NNFormer^37^ and UNETR^38^. The networks were trained by using the nnUNet pipeline, with a patch size of 128×128×128 voxels, channel-wise min-max normalization, initial learning rate of 0.001, learning rate decay, SDG optimizer, for 1000 epochs. We train using 5-fold cross validation, and evaluate voxel DICE, instance DICE^39^ and Betti Matching scores^40^. Based on two out of the three metrics, we select the 3D UNet for carrying out our downstream quantifications.

### Whole-body organ and tissue segmentation (Tissue-Module)

For the segmentation of internal organs, we used six annotated mice (from the CD68-EGFP line) to train five different networks: 3D UNet^34^, V-NET^35^, Attention UNET^36^, NNFormer^37^, Swin UNETR^41^. All the architectures were trained through the nnUNET^21^ pipeline using z-score normalization of each channel, and foreground oversampling. The networks were trained with Stochastic Gradient Descent (SGD) optimizer, using a batch size of 2, patch size of 64×256×128 voxels, initial learning rate of 0.01 and learning rate decay, for a total of 1000 epochs. The resulting networks were evaluated on the four UCHL1 mice. During training, we performed 5-fold cross validation, and the final predictions were made by ensembling the five resulting networks. We report voxel-wise Dice scores in Suppl. Table 4. We identified the 3D UNet as the best performing network architecture, with the following properties: 6 downsampling layers, 5 upsampling layers, 3×3×3 sized convolutional blocks and **a** maximum feature size of 320 in the bottleneck.

Second, we train a model to segment the soft tissues of mice, such as muscle and adipose tissue. We iteratively increased the size of our annotated dataset through inference on un-annotated patches, and manual correction of the wrongly segmented areas. As a result, our final networks were trained on a dataset of 387 samples containing a total volume of 2 billion voxels of adipose tissue, and 2 billion voxels of muscle. We then train on these patches the following neural network architectures: 3D UNet, V-NET, Attention UNET, and UNETR. We trained using 5-fold cross-validation, and for evaluation, we report and select based on the validation scores of the ensembles of the 5 resulting networks (Suppl. Table 5). The networks were trained with SGD optimizer, using a batch size of 2, patch size of 128×128×128 voxels, initial learning rate of 0.001, and learning rate decay, for a total of 1000 epochs. Again, the convolutional 3D UNET performs best among the implemented baselines.

The final inference pipeline for the Tissue-Module is carried out by sequential inference of our model ensemble for first, the organs, and then for the tissues. Specifically, first, the autofluorescence and PI channels of the acquired LSFM stack are downsampled to a resolution of 59×59×60 um/ voxel, and saved as a 3D Nifti volume. This is then fed into the organ segmentation network. The result is a 3D volume containing the masks of the 20 organs of interest, which can be used downstream for localizing structures of interest within organs, or for the quantification of organ volumes. Next, the organ masks are upsampled, and a “non-organ” mask is calculated, which is applied to the original scan. Through this process, we obtain a mask of the mouse volume that does not contain internal organs. As the size of the resulting data is too large to efficiently process, the “non-organ mouse images” are cropped into sub-volumes of 500 × 500 × 500 voxels, which are fed into the tissue segmentation network. These patches can then be reconstructed, resulting in a full-resolution tissue map. Lastly, by combining the organ maps and the tissue maps, we obtain a spatial segmentation of major organs and tissues in the mouse body.

### Whole-body inference of CD68-EGFP mice

For inference of CD68-EGFP on the whole-body mouse scans, we first cropped the autofluorescence and CD68 channels of the LSFM scans. For this, we used the same patch sizes and distribution as for the tissue segmentation. Then, a subset of the resulting sub volumes was selected, based on whether these were within the body of the mice. These patches were run through the CD68+ segmentation network. The resulting binary masks were split into components by using the cc3d library^29^ for connected component analysis. Subsequently, each individual detected connected component was post-processed by storing its location, volume, center of mass, and shape^27^. Next, we combined the inference results of our ensemble. Based on location of the center of mass, we automatically assign each segmented CD68-EGFP+ cluster to the internal organs or the segmented tissues, with blobs not located in any of these being discarded as false positives. We further discard components whose shape was elongated (string-like) as false positives, as these can often be artifacts, representing high-contrast blood vessels or nerves. Lastly, we grouped the detected CD68-EGFP clusters into three discrete categories, based on their volume (amount of segmented voxels within a component): small (smaller than 50 voxels), medium (between 50 and 500 voxels), and large (over 500 voxels). We chose these categories based on the observation that, when considering the total spatial volume of all clusters, each of these three categories would represent a similar amount (approximately 30%) of the total CD68+ segmented volume. Then, for each mouse and for each organ or tissue, we studied the % composition of each of these categories, and analyzed differences between the Chow and HFD groups.

While applying the CD68 segmentation network to the whole mouse bodies, we observe it displays zero-shot transfer learning abilities in the limited setting of applying the model in inference to certain novel tissues, where we observe positive detections. Hence, in order to validate any reported changes, we perform a) visual analysis of the resulting segmentation, and b) a VR-based annotation of a representative test patch in the tissue of interest. We compare the result of the automatic segmentation against the manual annotation in order to evaluate the network’s transfer learning abilities. We only consider valid quantifications where the network passes with a DICE score>65%.

### Whole-body inference of UCHL1-EGFP mice

To apply the nerve segmentation network to whole-body scans in full resolution efficiently, we adapted the sliding window inference method previously used for segmentation tasks in medical image (MONAI^42^) and the mouse brain (DELIVR^10^). Our inference is implemented using the highly efficient ZARR file format and DASK parallel computing framework, enabling lazy loading and multiprocessing for data handling and writing tasks and therefore a rapid full body analysis.

Before inference, we applied percentile normalization to each scan, similar to the model training stage. Given the significant imbalance between nerve voxels and background voxels in whole-body scans, we computed the 0.10th percentile and 99.9th percentile of all non-zero voxel intensity values to set the minimum and maximum thresholds, to effectively enhance the contrast between nerves and the background.

After inference, we obtained the whole-body nerve segmentation of UCHL1-EGFP mice. We then performed connected component analysis to post-process the segmentation results, eliminating large false positive segments caused by high-intensity regions within the mouse body. Subsequently, we quantified the nerve voxels and density from three perspectives: the entire body, individual tissues, and specific organs.

To quantify nerves in the entire body, the organ and tissue segmentations from the Tissue-Module were combined to form a binary mask of major organs and tissues in the mouse body. By dilating this binary mask, we created a whole-body mask that covers the entire mouse body, allowing us to compute the nerve voxels and density within. For tissue wise quantification, the tissue segmentation from the Tissue-Module was utilized to calculate the nerve voxels and density in fat and muscle tissues. For quantifying nerves for specific organs, we accounted for structures in the immediate vicinity of the organs by extending the organ segmentation by a 15-voxel boundary to calculate the organ wise statistics. Notably, to create the head mask, we overlaid the dilated brain masked with whole-body mask, resulting in a precise mask for quantifying the nerve voxels inside.

### Computational load of training and applying MouseMapper

The experiments presented in this work were carried out using a cohort of 12 mice (6 HFD feeding, 6 chow). Clearing and imaging these mice generated 46700 2D z-slices and 12 trillion voxels, occupying 10.35 Terabytes. To accurately quantify these data, our annotation efforts resulted in significantly ample datasets. For the Nerve-Module, we manually annotated 72 GB of data. While building the Immune-Module, we annotated 350 MB of data from representative areas in visceral and subcutaneous fat, as well as in the peritoneum. The organ segmenter of the Tissue Module was trained using 10 GB of downsampled organ data, whereas the tissue segmenter (for fat, muscle, bone and bone marrow) was trained using 46 GB of full-resolution tissue annotations, built as a mixture of manual and automatic annotations. In order to train the networks building our MouseMapper pipeline, as well as to run the predictions and quantifications presented in this paper, the High Performance Computing cluster of Helmholtz Zentrum Munich was used. Thus, the processes could be parallelized and carried out more efficiently. We estimate that a total of 7250 GPUhours were necessary to execute the presented experiments and quantifications.

### Graph extraction

Graph extraction was performed as previously described^11,43^. Similarly we extracted the skeletonization, depth map and extracted a graph of the resulting skeleton. All nodes with degree 2 were pruned from the graph, as well as small, isolated sub graphs. Since the resulting image data was too large to fit into a reasonable amount of RAM, we separated the whole image into sub blocks using dask. Then, we extracted the graphs from each sub block and merged them together. We fused all nodes together on the border between two blocks where the Euclidian distance between nodes was less than a given threshold by introducing a new edge between the nodes. We quantify the thickness of each node and each edge using the depth map, the degree of each node and the number of leaf nodes (nodes with degree = 1).

### Spatial proteomics sample preparation

For spatial proteomics of trigeminal ganglia of UCHL1-EGFP mice, 18G needle punches prepared from rehydrated trigeminal ganglia and subsequently used for proteomics sample preparations as described previously^7^. Briefly, the samples were resuspended in 6% SDS buffer, heat denatured at 95°C for 45 min at 600 rpm in a thermoshaker, sonicated in high mode for 30 cycles (30 sec OFF, 30 sec ON) (Bioruptor® Plus; Diagenode) and then precipitated using 80% acetone overnight in -20°C. The next day, these samples were centrifuged and the pellet was resuspended in SDC lysis buffer (2% SDC, 100 mM Tris-HCl pH 8.5). The samples in the SDC buffer were sonicated in high mode for 15 cycles (30 sec OFF, 30 sec ON) (Bioruptor® Plus; Diagenode). The samples were again heated at 95°C at 600 rpm in a thermoshaker for 45 min. The protein samples were digested with Trypsin and LysC (1:50, protease:protein ratio) at 37°C, 1,000 rpm shaking, overnight. Resulting peptides were acidified with 1% TFA 99% isopropanol with 1:1 volume-to-volume ratio, vortexed and centrifuged to pellet residual particles. The supernatant was transferred to fresh tubes and subjected to in-house built StageTip clean-up consisted of three layers of styrene divinylbenzene reversed-phase sulfonate (SDB-RPS; 3 M Empore) membranes. Peptides were loaded on the activated (100% ACN, 1% TFA in 30% Methanol, 0.2% TFA, respectively) StageTips, run through the SDB-RPS membranes, and washed by EtOAc including 1% TFA, isopropanol including 1% TFA, and 0.2% TFA, respectively. Peptides were then eluted from the membranes via 60 μL elution buffer (80% ACN, 1.25% NH4OH) and dried using a vacuum centrifuge (40 min at 45°C). Finally, peptides were reconstituted in 8-10 μL of loading buffer (2% ACN, 0.1% TFA) and stored in -80°C until further use.

### Liquid chromatography and mass spectrometry (LC-MS/MS)

The mass spectrometry data was acquired in data-independent acquisition (DIA) mode. The LC-MS/MS analysis was carried out using EASY nanoLC 1200 (Thermo Fisher Scientific) coupled with trapped ion mobility spectrometry quadrupole time-of-flight single cell proteomics mass spectrometer (timsTOF SCP, Bruker Daltonik GmbH, Germany) via a CaptiveSpray nano-electrospray ion source. Peptides (50 ng) were loaded onto a 25 cm Aurora Series UHPLC column with CaptiveSpray insert (75 μm ID, 1.6 μm C18) at 50°C and separated using a 50 min gradient (5-20% buffer B in 30 min, 20-29% buffer B in 9 min, 29-45% in 6 min, 45-95% in 5 min, wash with 95% buffer B for 5 min, 95-5% buffer B in 5 min) at a flow rate of 300 nL/min. Buffer A and B were water with 0.1 vol% formic acid and 80:20:0.1 vol% ACN:water: formic acid, respectively. MS data were acquired in single-shot library-free DIA mode and the timsTOF SCP was operated in DIA/parallel accumulation serial fragmentation (PASEF) using the high sensitivity detection-low sample amount mode. The ion accumulation and ramp time were set to 100 ms each to achieve nearly 100% duty cycle. The collision energy was ramped linearly as a function of the mobility from 59 eV at 1/K0 = 1.6 Vs cm^−2^ to 20 eV at 1/K0 = 0.6 Vs cm^−2^. The isolation windows were defined as 24 × 25 Th from m/z 400 to 1000.

### Proteomics data processing

diaPASEF raw files were searched against the mouse uniport database using DIA-NN (Ref. PMID: 31768060). Peptides length range from seven amino acids were considered for the search including N-terminal acetylation. Oxidation of methionine was set as a variable modification and cysteine carbamidomethylation as fixed modification. Enzyme specificity was set to Trypsin/P with 2 missed cleavages. The FASTA digest for library-free search was enabled for predicting the library generation. The FDR was set to 1% at precursor and global protein level. Match-between-runs (MBR) feature was enabled and quantification mode was set to “Robust LC (high precision)”. The Protein Group column in DIA-NN’s report was used to identify the protein group and PG.MaxLFQ was used to calculate the differential expression.

### Proteomics data analysis

Data were analyzed using scanpy (v. 1.10.1) and anndata (v. 0.8.0) in Python 3.10. Twelve independent samples were analyzed from each group (High-Fat Diet and Chow) from three animals with samples from both right and left trigeminal ganglia. All proteins expressed in less than half of the samples in each group were filtered out, resulting in 6686 proteins used for downstream analyses. The data was log-transformed and normalized per sample. The missing values were input using KNNImputer (n_neighbors=5) from sklearn package (v. 1.2.1). With scanpy’s dendrogram function scipy’s hierarchical linkage clustering was calculated on a Pearson correlation matrix over groups which was calculated for 50 averaged principal components. To identify differentially regulated proteins across two groups (HFD and Chow), we combined samples from the right and the left trigeminal ganglia. Differential expression analysis was conducted using Scanpy’s method ’ rank_genes_groups’ with method set to ’t-test We applied a threshold of p < 0.05 and |log fold change| > 0.5 to identify differentially expressed proteins (DEPs). These DEPs were subsequently visualized using volcano plots. Pathway enrichment analysis was performed on the combined up- and down-regulated proteins using the KEGG and Reactome databases. The most relevant pathways were highlighted, displaying the DEPs involved in each pathway.

## Statistical analysis

Results from biological replicates were expressed as mean ± s.e.m. Statistical analysis was performed using GraphPad Prism (v.9). To compare two conditions, unpaired Student’s t-tests or Mann–Whitney U-tests were performed. Insulin tolerance tests were analyzed using two-way ANOVA with Šídák’s multiple comparisons test. Proteomics data analysis was performed as described above.

## Data and Code Availability

Supplementary Videos can be seen here: http://discotechnologies.org/MouseMapper/

Whole body scans can be found to scroll through : [Chow-Nerve](https://wilddisco.translatum.xyz/datasets/erturk_lab/UCHL-1%20Chow%20778/view#3076,7390,1899,0,1.3), [HFD-Nerve](https://wilddisco.translatum.xyz/datasets/erturk_lab/UCHL1_HFD_788/view#3718,7491,1848,0,1.3), [Chow-Inflammation](https://wilddisco.translatum.xyz/datasets/erturk_lab/CD68_chow_7790/view#2545,7331,1679,0,1.3), [HFD-Inflammation](https://wilddisco.translatum.xyz/datasets/erturk_lab/CD68_HFD_7788/view#3429,7345,1926,0,1.3). Supplementary Videos can be viewed here:discotechnologies.org/MouseMapper.Our code will be made available here: https://github.com/erturklab/mouseMapper

## ADDITIONAL INFORMATION

### Author contributions

DK performed in vivo studies, vDISCO clearing, light-sheet imaging, data-processing, 3D-reconstructions, proteomic sample isolation, protein validations, data analysis, visualizations and project management. IH performed deep-learning analysis for immune cell segmentation, 3D tissue mapping, whole-body nerve inferences, data analysis and visualizations. RA performed deep-learning analysis for nerve segmentation, graph extraction, data analysis, visualizations and generated online atlases. ZIK supported in vivo experiments, performed vDISCO clearing, light-sheet imaging, data processing, data analysis and visualizations. YC performed deep-learning analysis of nerves, whole-body nerve inferences, data analysis and visualizations. LH, CC, VMK, NG performed VR-annotation. LH performed data visualizations. MT supported data processing and performed proteomics sample isolation. SK performed proteomics sample preparation, MS and data processing. MA analyzed proteomics data and performed visualizations. FK supported online atlas generation. PM, JG, DJ supported in vivo experiments. DJ performed the whisker stimulation. MP, DR, MR, BM, SH, MBD and AE provided funding. ME and FH provided critical input. MR, FH, ME, JCP, MBD edited the manuscript. HSB supported proteomics and data curation. SH and MBD provided resources. JCP provided critical input to the deep learning analysis and the manuscript. DK and AE conceptualized the study and drafted the manuscript. DK, IH, IK, ME and AE revised the manuscript. AE supervised and managed the entire project.

## Acknowledgements

We thank Lisa Mehr and Daniela Haß for excellent technical support. This work was supported by the Vascular Dementia Research Foundation, Deutsche Forschungsgemeinschaft (DFG, German Research Foundation) under Germany’s Excellence Strategy within the framework of the Munich Cluster for Systems Neurology (EXC 2145 SyNergy, grant no. ID 390857198) and DFG (grant nos. SFB 1052, project A9; TR 296 project 03) as well as the German Federal Ministry of Education and Research (Bundesministerium für Bildung und Forschung) within the NATON collaboration (grant no. 01KX2121) and the HIVacToGC collaboration. This work was also supported by the European Research Council Consolidator grant (CALVARIA, grant no. GA 865323 to A.E.) and Nomis Heart Atlas Project Grant (Nomis Foundation). This work was supported by the European Research Council under the European Union’s Horizon 2020 research and innovation program (949017 to M.R.) and a grant from the Else-Kröner-Fresenius-Stiftung (2020 EKSE.23 to S.H.), as well as the Edith-Haberland-Wagner Stiftung.

## Video legends

**Supplementary Video 1**. Whole-body reconstructions of normal (left) and obese (right) UCHL1-EGFP mice. Peripheral nerves (UCHL1-EGFP+) are shown in green, bones and organs are shown in red (propidium iodide labeled) and muscle (autofluorescence) is shown in blue.

**Supplementary Video 2**. Whole-body reconstructions of normal (left) and obese (right) CD68-EGFP mice. Immune cells (CD68-EGFP+) are shown in cyan, bones and organs are shown in magenta (propidium iodide labeled) and muscle (autofluorescence) is shown in yellow.

**Supplementary Video 3**. Whole-body reconstructions of a chow-fed CD68-EGFP mouse showing CD68-EGFP+ cells in the whole mouse. Immune cells (CD68-EGFP+) are shown in cyan, bones and organs are shown in magenta (propidium iodide labeled) and muscle (autofluorescence) is shown in yellow.

**Supplementary Video 4**. Whole-body reconstructions of a HFD-fed CD68-EGFP mouse showing infiltrating CD68-EGFP+ cells in the whole mouse with accumulations in ScAT, ViscAT and the peritoneum. Immune cells (CD68-EGFP+) are shown in cyan, bones and organs are shown in magenta (propidium iodide labeled) and muscle (autofluorescence) is shown in yellow.

**Supplementary Video 5**. Representative chow-fed mouse showing AI-segmented organs and tissue. Each color represents a different organ or tissue segmented using the Tissue-Module of MouseMapper.

**Supplementary Video 6**. Representative obese mouse showing AI-segmented organs and tissue. Each color represents a different organ or tissue segmented using the Tissue-Module of MouseMapper.

**Supplementary Video 7**. Representative obese CD68-EGFP mouse showing AI-segmented immune cell clusters (blue: small clusters, green: medium-sized clusters, red: large-sized clusters).

**Suppl. Figure 1:**
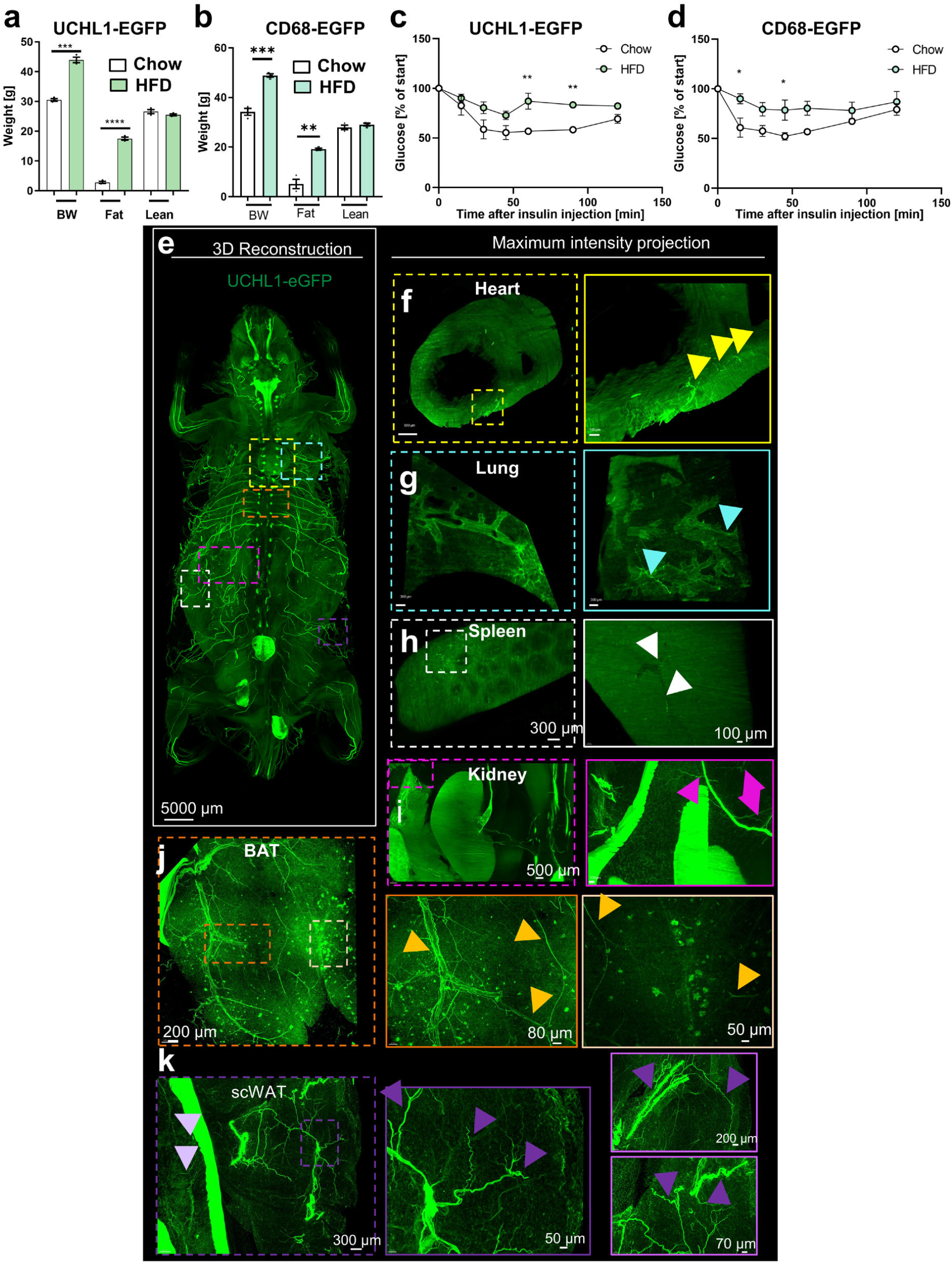
HFD-feeding associated with increased body weight. **a-b**, Body composition analysis using EchoMRI of UCHL1 and CD68 –EGFP mice, (n=3/group, BW: Body weight). **c-d** Insulin tolerance test. (n=3/group). *p<0.05, **p<0.01, ***p<0.001, ****p<0.0001. **e**, Representative 3D reconstruction of a HFD-fed UCHL1-EGFP animal to showcase expression of UCHL1-EGFP+ nerves shown in green in all panels. **e-k**, Zoomed in views of dashed regions in (f) showing the heart demonstrating the nerves in the heart with yellow arrows, lungs in (g) nerves with blue arrows, spleen in (h), nerves with white arrows, abdominal cavity with the kidneys and a part of the spleen in (i), nerves in the visceral fat are shown here with pink arrows, brown adipose tissue (BAT) in (j) different zoomed in regions show the innervation with orange arrows and lastly the the visceral and subcutaneous white adipose tissue (scWAT) in (k), light purple arrows show the visceral fat innervation and the zoomed images with dark purple outline and arrows show the subcutaneous adipose tissue nerve innervation. Right two panels depict different depths around the same subcutaneous fat region. The images are acquired with a 4x objective and are shown using different brightness and contrast settings in different organs. The thickness of the MIPs are as follows : 500 μm in (f), 100 μm in (g), 230 μm for left and 500 μm for right panel in (h), 1500 μm in (i), 600 μm in (j) and 1500 μm in (k).

**Suppl. Figure 2:**
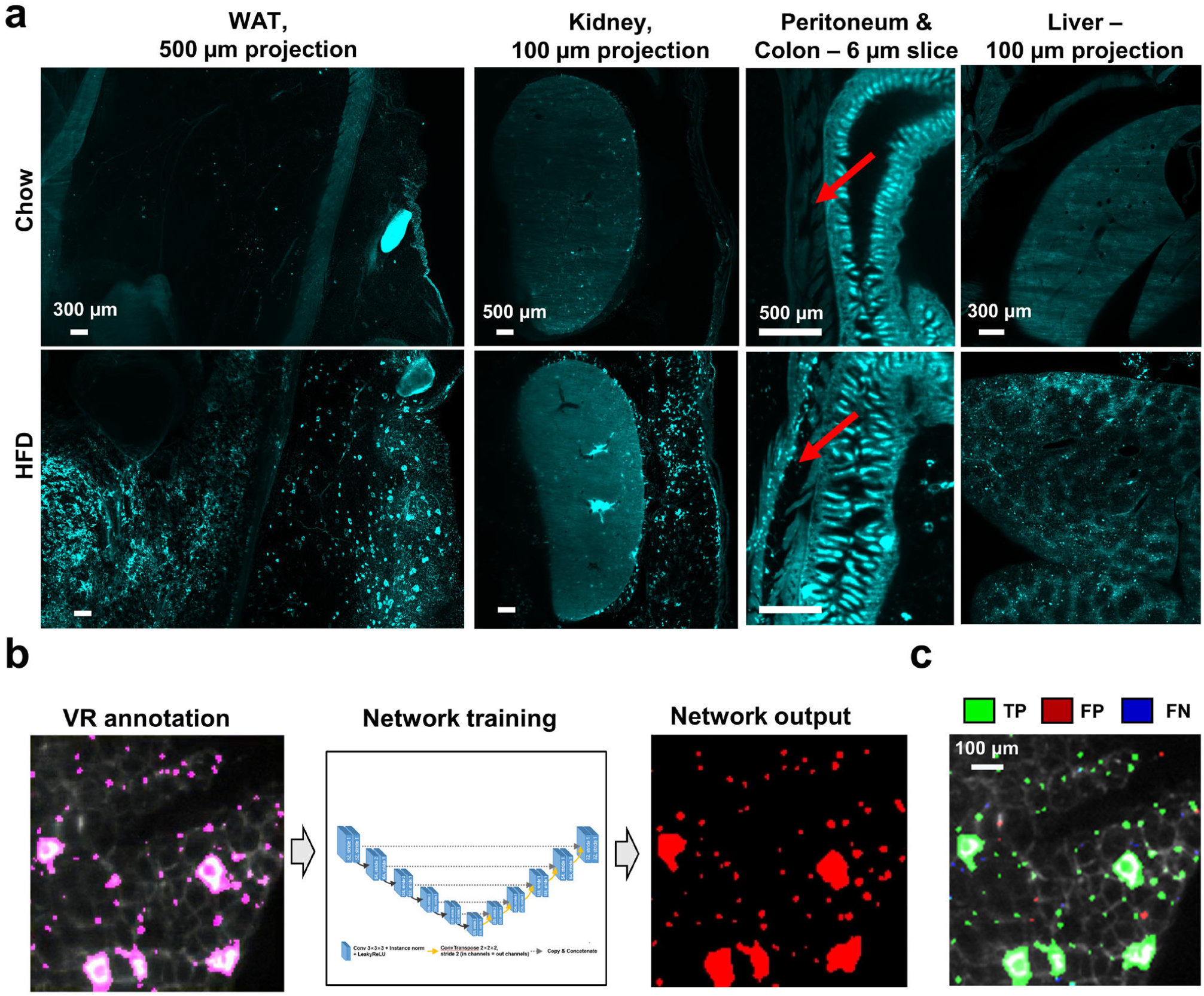
Development of the Immune-Module of MouseMapper. **a**, Images of indicated areas in chow and HFD-fed mice showing CD68-EGFP+ cells in cyan (WAT: white adipose tissue). **b**, To train the Immune-Module, CD68-EGFP+ cells were annotated using virtual reality (VR). Annotations were used to train a deep neural network, which generates the segmentation of CD68-EGFP+ cells as network output. **c**, 3D qualitative evaluation of the network performance for the segmentation of CD68-EGFP+ cells based on instance dice. Areas that overlap with reference annotations (TP) are masked in green, areas with no overlap in reference annotations (FP) are masked in red. Undetected reference annotation areas (FN) are marked in blue. TP, true positive; FP, false positive; FN, false negative.

**Suppl. Figure 3:**
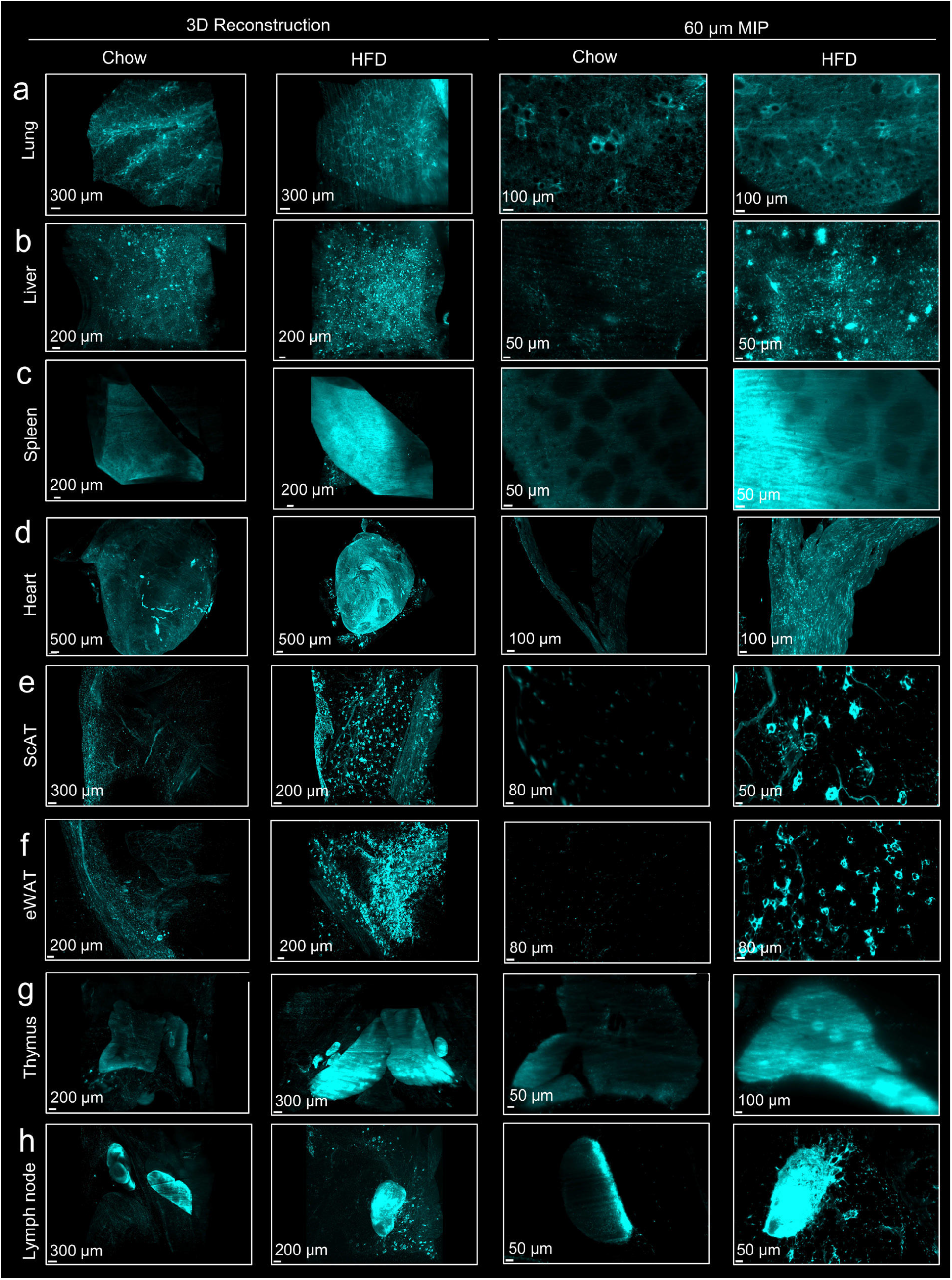
High resolution CD68-EGFP organ comparison. **a-h**, Representative 3D reconstruction and 2D 60 μm thick maximum intensity projections (MIP) of (**a**) lung, (**b**) liver, (**c**) spleen, (**d**) heart, (**e**) ScAT, (**f**) eWAT, (**g**) thymus and (**h**) accessory axillary lymph node from the CD68-eGFP mouse line after vDISCO and light sheet imaging for chow and high fat diet fed animals. The left two columns represent the 3D reconstruction of the organs and the right two columns represent the MIPs. For each representation chow fed animals are shown on the left column and the high-fat diet fed animals are shown in the right column. The images are acquired with a 4x objective and are visualized using the same brightness and contrast settings.

**Suppl. Figure 4:**
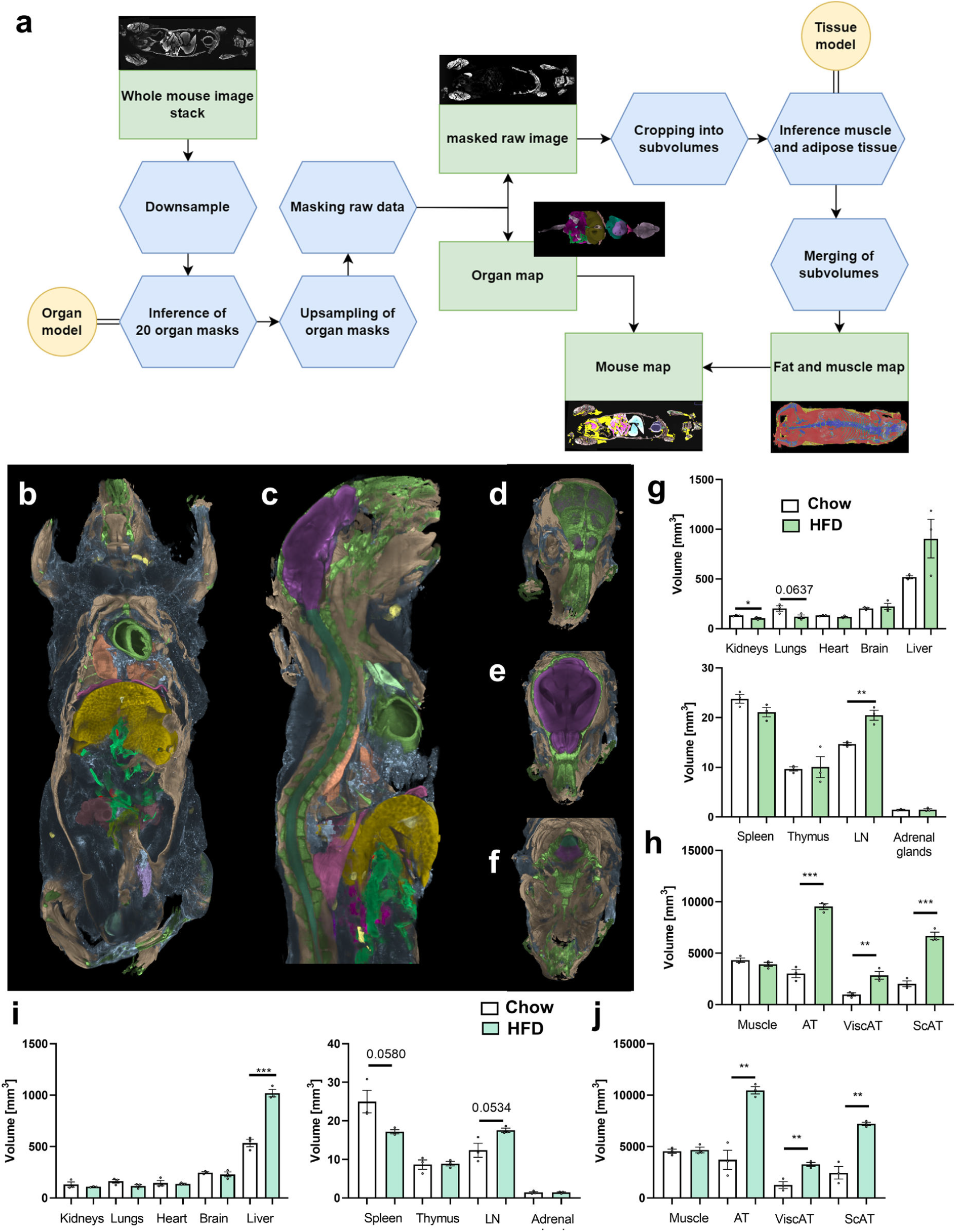
AI-based segmentation of organs and tissues with using the Tissue-Module of MouseMapper. **a**, Pipeline depicting the workflow of the organ and tissue segmentation model that was used in CD68-EGFP and UCHL1-EGFP mouse lines. **b-c**, Representative obese mouse showing AI-segmented organs and tissue. Each color represents a different organ or tissue segmented displaying b, ventral, c, sagittal view of the body. **d-f** Head from a representative chow animal in rostral view with AI-segmented organs and tissues in different z-planes. **g-h**, Organ (g) and tissue volumes (h) segmented in UCHL1-EGFP mice with the pipeline shown in **a**. White represents the chow group and green represents the HFD group. **i-j**, Organ (i) and tissue (j) volumes segmented in CD68-EGFP mice with the pipeline shown in a. White represents the chow group and cyan represents the HFD group. n=3/group, *p<0.05, **p<0.01, ***p<0.001.

**Suppl. Figure 5:**
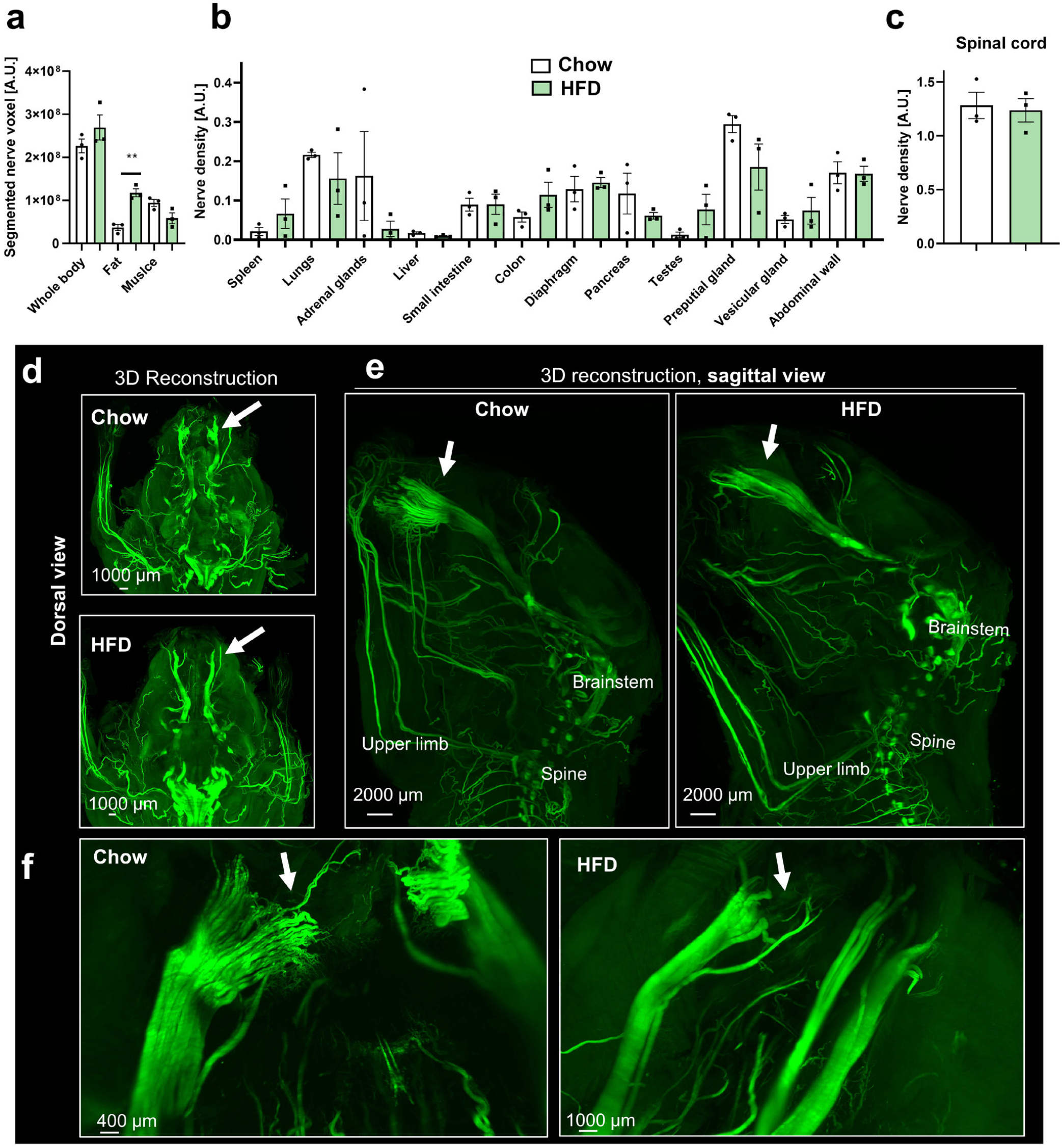
Quantification of segmented nerves and 3D visualization of UCHL1-EGFP mouse heads after vDISCO clearing, imaging, and reconstruction. **a**, Total nerve voxels of segmented nerves in the whole body, fat and muscle. **b-c**, Organ-wise quantification of nerve density in indicated organs/tissues. n=3/group, **p<0.01. **d**, Dorsal view of 3D reconstructions of heads of chow- and HFD-fed mice showing UCHL1-EGFP+ nerves. **e**, Sagittal view of 3D reconstructions of heads of chow- and HFD-fed mice showing UCHL1-EGFP+ nerves. **f**, Zoomed-in images of the intricate nerve structures. White arrows indicate structural changes in the infraorbital nerve across all panels.

**Suppl. Figure 6.**
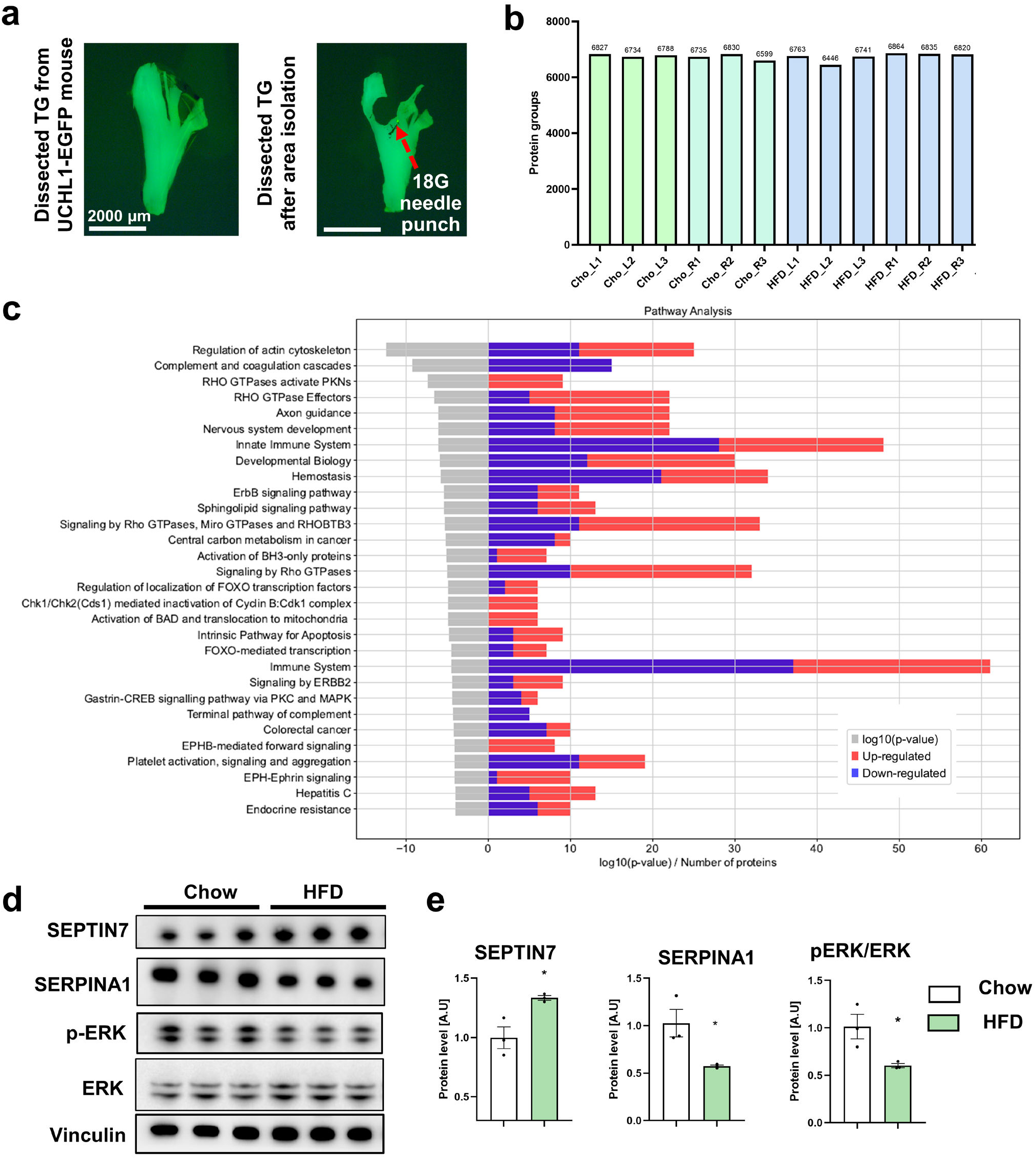
Trigeminal ganglion proteome differences in chow vs high-fat diet fed animals. **a**, Dissected trigeminal nerve from UCHL1-EGFP mouse before (left) and after (right) puncture with the 18G needle. Red arrow indicates the excised region that was subjected to proteomic analysis. **b**, number of protein groups detected in each proteomic sample. Cho stands for chow group and HFD stands for high-fat diet-fed group. L stands for the left trigeminal ganglion and R stands for the right trigeminal ganglion. **c**, Pathway analysis from the TG of chow vs HFD is depicted. The pathways that are affected are shown on the y-axis of the plot. Gray bar on the left side of the plot represents the log10 of p-value of each pathway whereas the right side of the plot depicts the number of proteins significantly different in each pathway, red representing the number of up-regulated proteins and blue representing the number of down-regulated proteins. **d**, Western blot of protein lysates from trigeminal ganglions of chow and HFD-fed mice to validate proteins that were differentially expressed in the proteomic analysis: SEPTIN7, SERPINA1, p-ERK, and ERK proteins using western blot. Vinculin is shown as representative loading control (n=3/group). **e**, Quantification of SEPTIN7, SERPINA1 and pERK/ERK protein levels detected in chow vs HFD groups (p<0.05 for each difference in protein expression, n=3/group).

**Supplementary Table 1:** Evaluation of different networks for nerve segmentation based on the volumetric dice. The best performing scores are highlighted in bold. Our final 3D Unet is composed of the same building blocks as the 3D Unet, but it has been trained with the addition of the Centerline DICE score to loss function and for an extended (2000) number of epochs.

|  | Attention Unet | NNFormer | SwinUNETR | UNETR | VNET | 3D Unet | Final 3D Unet |
| --- | --- | --- | --- | --- | --- | --- | --- |
| Voxel DICE Score | 0.6521 | 0.6205 | 0.6318 | 0.5180 | 0.5161 | **0.6687** | **0.7694** |

**Supplementary Table 2:** Evaluation of CD68 marker segmentation network. We report the average voxel and instance DICE scores, as well as Betti Matching score, obtained on the validation set at the end of training. Best scores are highlighted in bold. A high DICE score and low Betti Matching score indicate good performance.

| Score type\Network Architecture | AttentionUnet | NNFormer | UNETR | VNET | 3D UNet |
| --- | --- | --- | --- | --- | --- |
| Voxel DICE | 0.6685656 | 0.6895897 | 0.6835672 | **0.6987444** | 0.6868189 |
| Instance DICE | 0.83393675 | 0.8507939 | 0.8338425 | 0.8473859 | **0.85342973** |
| Betti Matching Score | 0.416529168 | 0.397961519 | 0.463957805 | 0.430044453 | **0.36935299** |

**Supplementary Table 3:** Transfer learning abilities of the CD68-EGFP segmentation network on new tissue types.

| Organ | Instance DICE score |
| --- | --- |
| Stomach | 0.6963 |
| Colon | 0.6832 |
| Liver | 0.6759 |

**Supplementary Table 4:** Organ segmentation performance of different baselines, per organ. We evaluated state of the art architectures for 3D structure segmentation on a stand-alone test set, comprised of 4 whole body mice (2 HFD, 2 chow). We report the average DICE score, per organ. Best performing scores for each case are highlighted in bold.

| Organ\<br>Network<br>architecture | AttentionUnet | NNFormer | SwinUNETR | 3D UNet |
| --- | --- | --- | --- | --- |
| Spleen | 0.91371775 | **0.94894576** | 0.93309945 | 0.91092151 |
| Kidneys | 0.97267586 | 0.93032014 | 0.89774877 | **0.9762361** |
| Lungs | 0.9401775 | 0.91957378 | 0.83477843 | **0.94498158** |
| Heart | **0.96156639** | 0.91270357 | 0.90297139 | 0.95824647 |
| Brain | 0.98283041 | 0.9677701 | 0.97718501 | **0.98289859** |
| Gallbladder | **0.7923587** | 0.49352396 | 0.54942632 | 0.7797811 |
| Adrenal glands | 0.9251489 | 0.82583666 | 0.79534006 | **0.93445438** |
| Liver | 0.95344108 | 0.93522882 | 0.91427165 | **0.95384806** |
| Thymus | 0.89085966 | 0.88994277 | 0.88707769 | **0.89394242** |
| Lymph nodes | **0.91784495** | 0.86091363 | 0.87262303 | 0.91692603 |
| Gut | 0.47927877 | 0.44504771 | 0.44276318 | **0.50523841** |
| Diaphragm | 0.72079068 | 0.70729929 | 0.69553173 | **0.74165958** |
| Pancreas | 0.58869672 | 0.56822401 | 0.41258866 | **0.62522465** |
| Abdominal wall<br>(peritoneum) | 0.84319705 | 0.82153213 | 0.82839638 | **0.84549576** |
| Spinal cord | 0.84208506 | 0.82547545 | 0.84191978 | **0.84481466** |
| Testes | **0.93766016** | 0.84609985 | 0.88627297 | 0.93748099 |
| Preputial gland | 0.92438507 | 0.90678233 | 0.88193274 | **0.92707711** |
| Peyer patches | **0.5689556** | 0.10222362 | 0.48704222 | 0.56012666 |
| Vesicular gland | **0.86798155** | 0.87190825 | 0.8211478 | 0.86536998 |
| Bladder | 0.49881184 | 0.33731034 | 0.35585785 | **0.51761812** |
| Average | 0.82612312 | 0.75583309 | 0.76089871 | **0.83111715** |

**Supplementary Table 5:** Tissue segmentation performance of different architectures for the indicated tissue types. We report the DICE score of 5-fold cross validation result on the final epoch. Best performance is highlighted in bold.

| Tissue\<br>Network<br>architecture | AttentionUnet | NNFormer | UNETR | VNET | 3D UNet |
| --- | --- | --- | --- | --- | --- |
| Fat | 0.904±0.017 | 0.864±0.038 | 0.864±0.032 | 0.152±0.304 | **0.908±0.021** |
| Muscle | 0.958±0.014 | 0.933±0.013 | 0.941±0.015 | 0.157±0.314 | **0.961±0.009** |
| Bone | 0.740±0.061 | 0.669±0.046 | 0.671±0.059 | 0.102±0.203 | **0.756±0.046** |
| Bone<br>Marrow | 0.881±0.055 | 0.853±0.037 | 0.865±0.018 | 0.129±0.258 | **0.887±0.028** |
| Mean±STD | 0.873±0.028 | 0.832±0.018 | 0.839±0.020 | 0.224±0.228 | **0.880±0.017** |

**Supplementary Table 6:**
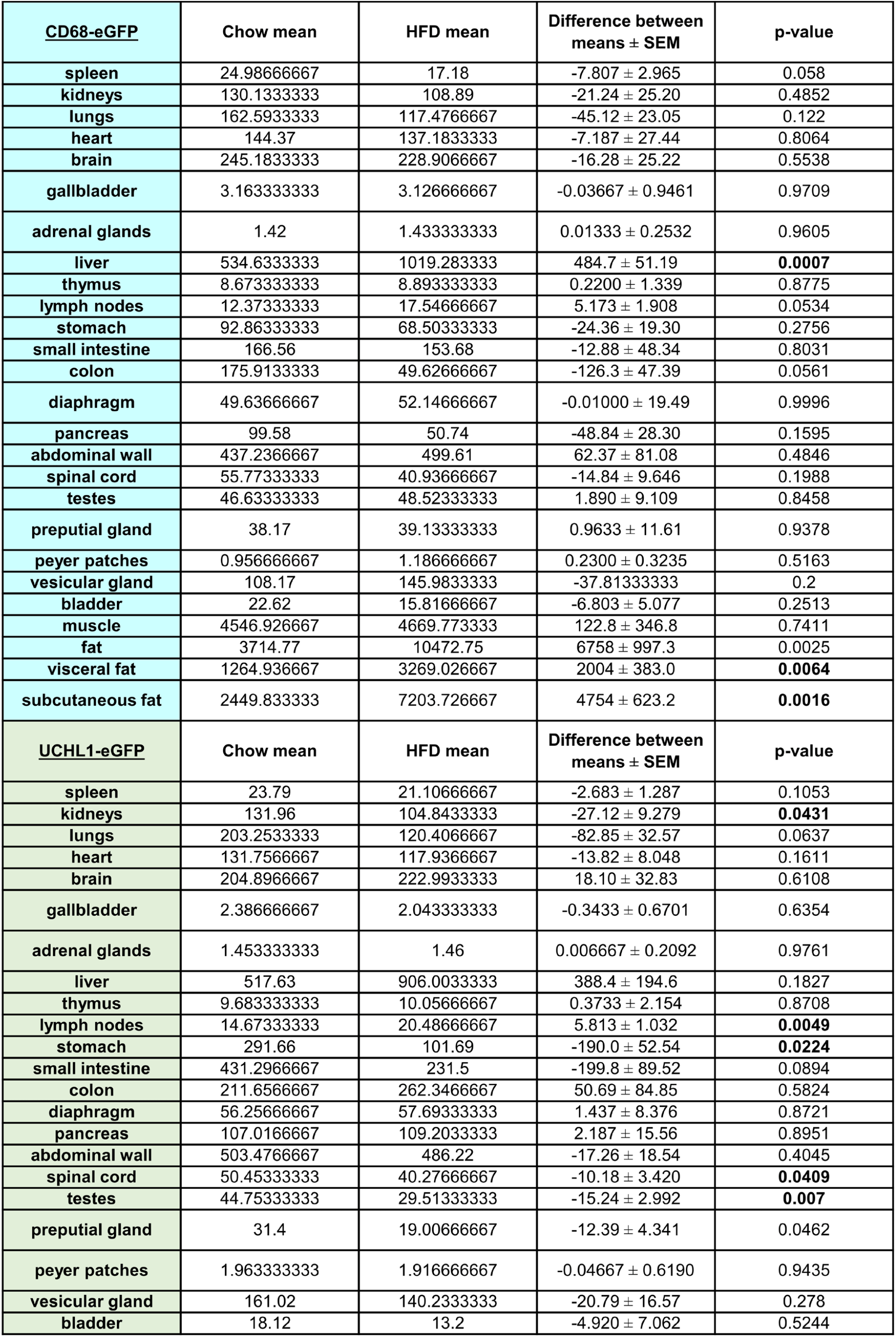
Organs and tissue volumes from CD68 and UCHL1-EGFP mouse scans were determined by Al-based segmentation.

